# The hepatic compensatory response to elevated systemic sulfide promotes diabetes

**DOI:** 10.1101/2020.04.27.064287

**Authors:** Roderick N. Carter, Matthew T.G. Gibbins, Martin E. Barrios-Llerena, Stephen E. Wilkie, Peter L. Freddolino, Marouane Libiad, Victor Vitvitsky, Barry Emerson, Thierry Le Bihan, Madara Brice, Huizhong Su, Scott G. Denham, Natalie Z.M. Homer, Clare Mc Fadden, Anne Tailleux, Nourdine Faresse, Thierry Sulpice, Francois Briand, Tom Gillingwater, Kyo Han Ahn, Subhankar Singha, Claire McMaster, Richard C. Hartley, Bart Staels, Gillian A. Gray, Andrew J. Finch, Colin Selman, Ruma Banerjee, Nicholas M. Morton

## Abstract

Impaired hepatic glucose and lipid metabolism are hallmarks of type–2 diabetes. Increased sulfide production from cysteine, or sulfide–donor compounds, may beneficially regulate hepatic metabolism. Disposal of sulfide through the sulfide oxidation pathway (SOP) is critical for maintaining sulfide within a safe physiological range. We show that mice lacking the liver–enriched mitochondrial SOP enzyme thiosulfate sulfur–transferase (*Tst^−/−^* mice) exhibit high circulating sulfide, increased gluconeogenesis, hypertriglyceridemia and fatty liver, despite whole–body insulin–sensitisation. Unexpectedly, hepatic sulfide levels were normal in *Tst^−/−^* mice, a result of homeostatic induction of mitochondrial sulfide disposal and glutathione excretion associated with net suppression of protein persulfidation and nuclear respiratory factor–2 target proteins. Proteomic and persulfidomic profiling converged on gluconeogenesis and hepatic lipid metabolism and revealed a selective deficit in medium–chain fatty acid oxidation in *Tst^−/−^* mice. We reveal a critical role for TST in hepatic metabolism that raises implications for sulfide-donor strategies in the context of liver function and metabolic disease.

## Introduction

The prevalence of Type 2 diabetes (T2D) continues to soar in parallel with that of obesity^1^. Increased hepatic glucose production and aberrant hepatic lipid metabolism are cardinal features of T2D^2,3^. Dysregulation of hepatic nutrient metabolism in T2D is a promising area for therapeutic intervention because it precipitates the more severe liver pathologies that manifest along the spectrum of non-alcoholic fatty liver disease (NAFLD), steatosis, steatohepatitis and hepatocellular carcinoma^4^.

Hydrogen sulfide (hereafter referred to as sulfide), an endogenously produced gaseous signalling molecule^5–8^, has recently emerged as a modulator of nutrient metabolism^9–12^. Enzymatic sulfide production from sulfur amino acids is catalysed by cystathionine beta–synthase; CBS, and cystathionine gamma lyase; CTH^13,14^ and by 3-mercaptopyruvate sulfur transferase; MPST^15–17^. Thioredoxin-mediated reduction of cysteine persulfides on proteins also regulates free sulfide and cysteine persulfide levels^18^. Endogenously produced and exogenously administered sulfide specifically influences hepatic glucose and lipid metabolism^19,20^. Thus, *in vitro*, treatment of murine hepatocytes with NaHS, or overexpression of rat *Cth* in HepG2 liver cells, increased glucose production through increased gluconeogenesis and reduced glycogen storage^21^. Conversely, glucose production was lower in hepatocytes from *Cth* gene knockout mice (*Cth^−/−^* mice) that exhibit low sulfide production^21^. Elevation of sulfide with NaHS administration *in vivo* reduced cholesterol and triglyceride accumulation in the liver of high fat diet (HFD)–fed mice^22^. In contrast, inter-cross of sulfide production–deficient *Cth*^−/−^ mice with the hyperlipidemic *Apoe^−/−^* mouse strain (*Cth^−/−^Apoe^−/−^* mice) produced a phenotype of elevated plasma cholesterol following exposure to an atherogenic diet^23^. Consistent with their higher cholesterol, *Cth^−/−^Apoe^−/−^* mice developed fatty streak lesions earlier than *Apoe^−/−^* mice, and this effect was reversed by NaHS administration^23^. Sulfide may also indirectly impact hepatic nutrient metabolism through its effect on hepatic artery vasorelaxation and thus liver perfusion^24,25^. The apparently beneficial effects of sulfide administration in multiple disease indications has led to a major drive towards development of targeted H_2_S–donor molecules as a therapeutic approach^26,27^. However, an often-overlooked aspect of net sulfide exposure, key to the efficacy of therapeutic H_2_S–donors, is that it is regulated through its oxidative disposal. Thus, endogenous sulfide exposure is actively limited to prevent mitochondrial respiratory toxicity^28–30^. Sulfide is rapidly oxidized^31,32^ through the mitochondrial sulfide oxidation pathway (SOP), consisting of sulfide quinone oxidoreductase (SQOR), persulfide dioxygenase (ETHE1/PDO) and thiosulfate sulfurtransferase (TST, also known as rhodanese)^31,33,34^. The liver is highly abundant in SOP enzymes, and is a major organ of whole body sulfide disposal^32^. Mice lacking the *Ethe1* gene (*Ethe1^−/−^* mice) die of fatal sulfide toxicity^30^, consistent with its critical role in sulfide oxidation and the severe pathological consequences of unchecked sulfide build-up in tissues. However, the importance of mitochondrial TST in the SOP *in vivo* remains obscure. In contrast to *Ethe1^−/−^* mice, *Tst^−/−^* mice were grossly normal despite exhibiting substantially elevated blood sulfide levels, as implied by qualitative measures^35^. This revealed an important yet distinct role for TST in the SOP for the first time *in vivo*. Nevertheless, *Tst^−/−^* mice showed an apparently diabetogenic impairment of glucose tolerance^35^, consistent with the concept that increased sulfide promotes hepatic glucose production^21^. As *Tst* deficiency represents a model of chronic but viable sulfide elevation, determining the molecular mechanisms driving their aberrant metabolic profile can provide important insights into the optimal range for therapeutic sulfide exposure, particularly in light of the current interest in developing mitochondrially–targeted sulfide–donors^36,37^. To this end we sought to define the impact of *Tst* deficiency on the underlying molecular pathways that impact hepatic metabolism.

## Results

### Tst^−/−^ mice exhibit increased hepatic gluconeogenesis and dyslipidaemia despite mild peripheral insulin sensitisation

TST *mRNA* expression is highest in the liver (e.g. it is ~8-fold higher than in white adipose tissue http://biogps.org/#goto=genereport&id=22117). We therefore hypothesised that liver TST deficiency was the principal driver of the impaired glucose tolerance previously observed in *Tst^−/−^* mice^35^. *Tst^−/−^* mice exhibited higher glucose levels than C57BL/6J controls in response to a pyruvate challenge, consistent with higher hepatic glucose production (Figure 1A). We next tested phosphoenolpyruvate carboxykinase (PEPCK) activity, a key enzyme of *de-novo* hepatic glucose synthesis, and found it was higher in liver homogenates from *Tst^−/−^* mice (Figure 1B). Next, we performed a 1 hour ^13^C_3_-pyruvate metabolite–pulse incorporation experiment in isolated hepatocytes cultured in ^12^C_3_-pyruvate–free medium. Hepatocytes from *Tst^−/−^* mice displayed ^13^C labelling consistent with increased metabolism of pyruvate to oxaloacetate – a critical early step in gluconeogenesis. Specifically, aspartate, which is derived from pyruvate via oxaloacetate was significantly increased in the *Tst^−/−^* (Figure 1C). A trend towards higher ^13^C_3_ malate, and lower ^13^C_2_ acetyl–CoA was also observed (Figure S1A and S1B). ^13^C_3_ Lactate was similar between genotypes, suggesting a similar activity of glycolytic disposal of pyruvate through lactate dehydrogenase (Figure S1A and S1B). Total pool sizes for all measured metabolites were similar between genotypes (Figure S1C). Consistent with increased endogenous glucose production in the *Tst^−/−^* mice, fasting plasma glucose was higher in *Tst^−/−^* mice relative to 6J mice during the pre-clamp 3-^3^H glucose tracer infusion phase (60-90 minutes post tracer) of euglycemic, hyperinsulinemic (EH) clamp experiments (Figure 1D, Table S1A). Higher plasma glucose levels in *Tst^−/−^* mice under these conditions was not explained by lower glucose utilization in *Tst^−/−^* mice; glycogen synthesis and glycolysis were comparable between genotypes across 60–90 minutes (Table S1A). Glucose turnover – a derived parameter used to infer glucose production – was also comparable between genotypes (Table S1A). However, derivation of glucose turnover requires that glucose levels are stable during the period in which it is calculated. In our pre-clamp baseline period, a highly significant effect of time (Figure 1D) indicated that this assumption was not met, and thus true endogenous glucose production cannot be inferred from the glucose turnover parameter in this instance. Combined with the pyruvate tolerance, PEPCK activity and ^13^C_3_-pyruvate pulse data, higher fasting glucose levels in the *Tst* ^/^ mice, given comparable glucose utilization, is most likely due to higher endogenous glucose production in the *Tst^−/−^* mice.

**Figure 1.**
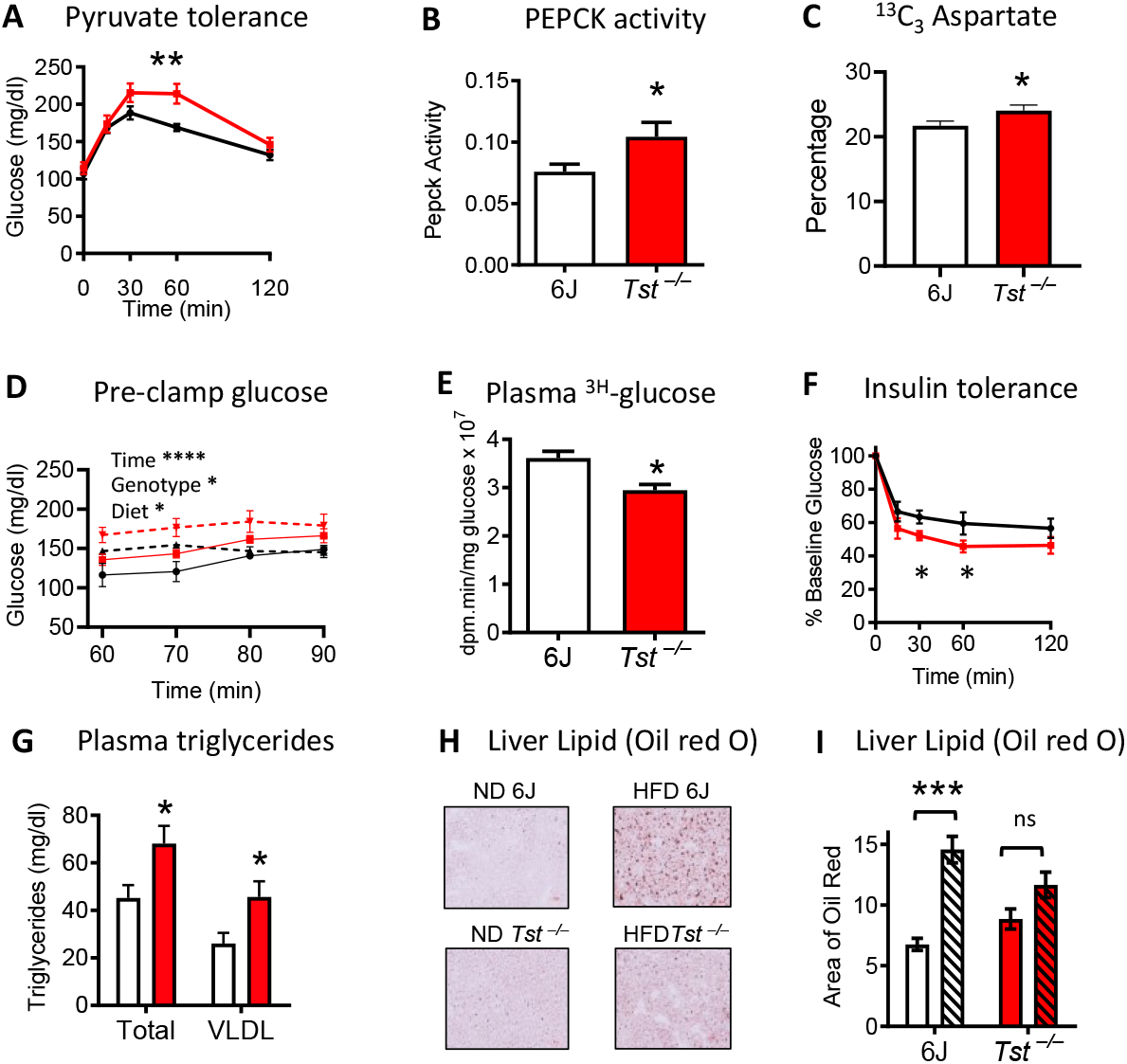
*Tst* deletion results in impaired glucose and lipid metabolism. (**A**) Plasma glucose over 120 minutes, following pyruvate (i.p., 1.5mg/g) administration in overnight fasted C57Bl/6J (black line, n = 9) and *Tst^−/−^* (red line, n = 8) normal diet-fed mice. (**B**) Extinction of NADH measured by absorbance at 340nm, coupled to PEPCK activity from liver homogenates taken from C57Bl/6J (white bar, n = 6) and *Tst^−/−^* (red bar, n = 6) normal diet-fed mice. (**C**) Production of ^13^C (M+2) acetyl-CoA and ^13^C (M+3) aspartate generated after a 1 hour pulse of 1mM 3-carbon labelled ^13^C (M+3) pyruvate in ^12^C pyruvate free media, expressed as a percentage of the total amount of detected metabolite, in primary hepatocytes from C57Bl/6J (white bars, n = 6) and *Tst^−/−^* (red bars, n = 5) normal diet-fed mice. (**D**) Blood glucose during the pre-clamp phase of the hyperinsulinemic, euglycemic clamp from C57Bl/6J (black lines), and *Tst^−/−^* (red lines) fed a control (ND, solid lines, n = 3, 6) or high fat diet (HFD, broken lines, n= 6, 7). (**E**) Mean integrated radioactive glucose (inversely related to whole body glucose uptake) during a hyperinsulinemic, euglycaemic clamp from normal diet fed C57Bl/6J control (white, n = 3), and *Tst^−/−^* (red, n = 6) mice. (**F**) Plasma glucose expressed as % of baseline glucose, over 120 minutes following insulin (i.p., 1mU/g) administration in 4 hour fasted C57Bl/6J (black line, n = 8) and *Tst^−/−^* (red line, n = 7) normal diet-fed mice. (**G**) HPLC quantified total and VLDL plasma triglyceride in 4 hour fasted C57Bl/6J (white bar, n = 6), and *Tst^−/−^* (red bar, n = 6) normal diet-fed mice. (**H**) Representative light microscopic images of fixed liver stained with Oil-Red O from normal diet-fed (ND) or high fat diet-fed (HFD) C57Bl/6J and *Tst^−/−^* mice. (**I**) Analysis of the area of red staining (Oil Red O) after thresholding, using Image J, from normal diet-fed (no pattern, n = 3-4/genotype) or high fat diet-fed (hatched pattern, n = 4-5/genotype) C57Bl/6J (white bars) and *Tst^−/−^* (red bars) mice. Data are represented as mean ±SEM. Significance was calculated using repeated measures ANOVA (**A**,**F**) 2-WAY ANOVA (**C**,**I**), 3-WAY repeated measures ANOVA (**D**) or unpaired two-tailed student’s t-test (**B**,**E**,F,**G**) * P < 0.05, ** P < 0.01, *** P < 0.001, **** P < 0.0001. For (**C**) the 2-WAY ANOVA revealed no overall effect of genotype, however a significant interaction between metabolite and genotype was found. The * on the histogram represents this interaction. For (**D**) significant effects of time (****), diet (*) and genotype (*) were found. For (**F**) the analysis was performed on absolute glucose values and demonstrated a significant effect of time (****) and an interaction between time and genotype (*). T-tests revealed that the decrement of glucose from baseline at 30 and 60 minutes after insulin was greater in the *Tst^−/−^* (*). For (**I**) no main genotype effect was found, but a significant effect of diet (***), and an interaction (*) were found. Post Hoc analysis using Sidaks’ multiple comparison test show an effect of diet on the 6J controls (***), whereas no effect of diet is found on the *Tst^−/−^*.

We next wished to explore whether the changes to glucose metabolism were driven by insulin resistance. Liver glycogen, a marker of long–term carbohydrate storage typically impaired with insulin resistance, was comparable between *Tst^−/−^* and C57BL/6J control mice (Figure S2A). Despite unchanged steady–state markers of hepatic insulin sensitivity, impaired glucose tolerance, previously described in the *Tst^−/−^* mice^35^, suggested that whole body, and usually hepatic, insulin–resistance was present. We investigated this using the euglycaemic clamp where, unexpectedly, we observed whole–body insulin sensitisation under these short–term steady–state conditions. During the clamp, when insulin was high, and blood glucose levels were maintained constant, the glucose infusion rate was comparable between genotypes (Table S1B). However, an increase in whole–body glucose uptake (integral glucose) by tissues in the *Tst^−/−^* mice was apparent (Figure 1E, Table S1B), supporting increased peripheral insulin–sensitivity, with a directionally consistent trend for increased glucose uptake into several tissues. We confirmed this finding using standard insulin tolerance tests, where the glucose decrement in response to insulin was greater in *Tst^−/−^* mice (Figure 1F, Figure S2B). Together these data demonstrate a net increase in dynamic whole–body insulin sensitivity, despite increased hepatic glucose output in *Tst^−/−^* mice. Finally, we assessed whole–body glucose homeostasis with the EH–clamp method after chronic HFD feeding. Under these conditions, *Tst^−/−^* mice maintained increased heptic glucose output (Figure 1D) but showed convergence of the insulin–sensitivity profile to that of the C57BL/6J mice.

We also assessed whether *Tst* deficiency was associated with impaired lipid metabolism, another hallmark of diabetes. Fast protein liquid chromatography analysis of triglyceride levels and their lipoprotein distribution revealed significantly higher total plasma triglycerides in *Tst^−/−^* mice (Figure 1G). The higher triglyceride was selectively associated with an increased VLDL triglyceride fraction (Figure 1G), consistent with a dominant liver-driven impairment in lipid metabolism^38^. Total and distinct lipoprotein fraction plasma cholesterol levels were similar between genotypes (Figure S2C and S2D), suggestive of a triglyceride–selective effect of *Tst* deficiency on hepatic lipid efflux. HFD significantly increased liver lipid content of C57BL/6J mice but did not further increase the elevated lipid levels in the liver of *Tst^−/−^* mice (Figure 1H-1I).

### TST deficiency elicits compensatory hepatic sulfide disposal mechanisms that drive reduced global protein persulfidation

A role for TST in the disposal of sulfide was suggested by its participation in the SOP^31,34^ and supported *in vivo* by the qualitatively higher blood sulfide of *Tst^−/−^* mice^35^. Here, we quantified circulating sulfide, showing approximately 10–fold elevation in the blood and plasma of *Tst^−/−^* mice (Table 1A and 1B). Thiosulfate, an oxidised metabolite of sulfide ^39,40^ and a TST substrate^41^, was approximately 20–fold higher in plasma (Table 1B), and profoundly higher (450–fold) in urine (Table 1C) of *Tst^−/−^* mice compared to C57Bl/6J mice. Reduced glutathione (GSH) levels were ^~^2–fold higher in the plasma of *Tst^−/−^* mice (Table 1B). To determine any direct hepatic contribution to the elevated systemic sulfide *in vivo*, whole blood was sampled from the inferior vena cava (IVC) (Table 1D). IVC sulfide levels tended to be higher in the *Tst^−/−^* mice, but the magnitude of the increase (~3-fold) did not parallel that in trunk blood (~10-fold), suggesting liver was not a major source of the elevated circulating sulfide. Surprisingly, liver homogenate sulfide, thiosulfate, cysteine and GSH levels were similar between *Tst^−/−^* and C57BL/6J mice (Table 1E). Further, cultured hepatocytes from *Tst^−/−^* and C57BL/6J mice exhibited similar intracellular sulfide levels, as estimated using P3, a sulfide–selective fluorescent probe^42^ (Table 1F). Mitochondrial sulfide levels in liver reported by MitoA/MitoN^43^ were similarly unchanged between genotypes (Table 1G). The apparently unaltered hepatic steady-state sulfide levels, despite higher circulating sulfide, suggested a profound homeostatic mechanism was invoked in the liver of *Tst^−/−^* mice. We assessed respiratory sulfide disposal (antimycin–sensitive) and found this was markedly increased in hepatocytes from *Tst^−/−^* mice, whereas antimycin–insensitive sulfide disposal was relatively reduced compared to hepatocytes from C57BL/6J mice (Table S2). Isolated liver mitochondria from *Tst^−/−^* hepatocytes also exhibited a higher sulfide disposal rate (Table S2). In addition, cysteine and GSH was excreted at higher levels from *Tst^−/−^* hepatocytes under basal conditions and after stimulation of sulfur amino acid metabolism by addition of methionine (Figure 2A and 2B). Consistent with higher GSH turnover, hepatocytes from *Tst^−/−^* mice showed resistance to exogenous H_2_O_2_–mediated mitochondrial ROS production (Figure S3). We next determined the global hepatic protein persulfidation profile, the major post–translational modification mediated by sulfide^44–46^. Mass spectrometry analysis of maleimide–labelled liver peptides revealed a greater abundance of peptides with a lower persulfidation level (under-persulfidated) in the liver of *Tst^−/−^* mice (Figure 2C). We confirmed this using semi-quantitative western–blot analysis on pulled down maleimide–labelled proteins (Figure 2D). Gene Ontology (GO) analysis of proteins exhibiting altered persulfidation levels revealed 28 terms different between *Tst^−/−^* and C57BL/6J mice by log–fold change (Table 2). Major under-persulfidated terms (20) included *“FAD-binding, methyl transferase, peroxisome, acyl-CoA dehydrogenase activity and transaminase”* while the less abundant over-persulfidated terms (8) were dominated by terms related to *“Nicotinamide metabolism”*. Given evidence for altered gluconeogenesis above, we also performed a pathway specifc analysis of the persulifidome. Persulfidation of proteins involved in gluconeogenesis showed a mixture of under and over–persulfidated cysteines with a strong enrichment of over-persulfidation, a pattern that was significantly different from the overall persulfidation profile of the liver from *Tst^−/−^* mice (Figure S4A). We further found that the gluconeogenesis pathway also exhibited a significantly higher magnitude of change (independent of direction of change) in persulfidation than in the overall profile of the liver of *Tst^−/−^* mice (Figure S4B).

**Figure 2.**
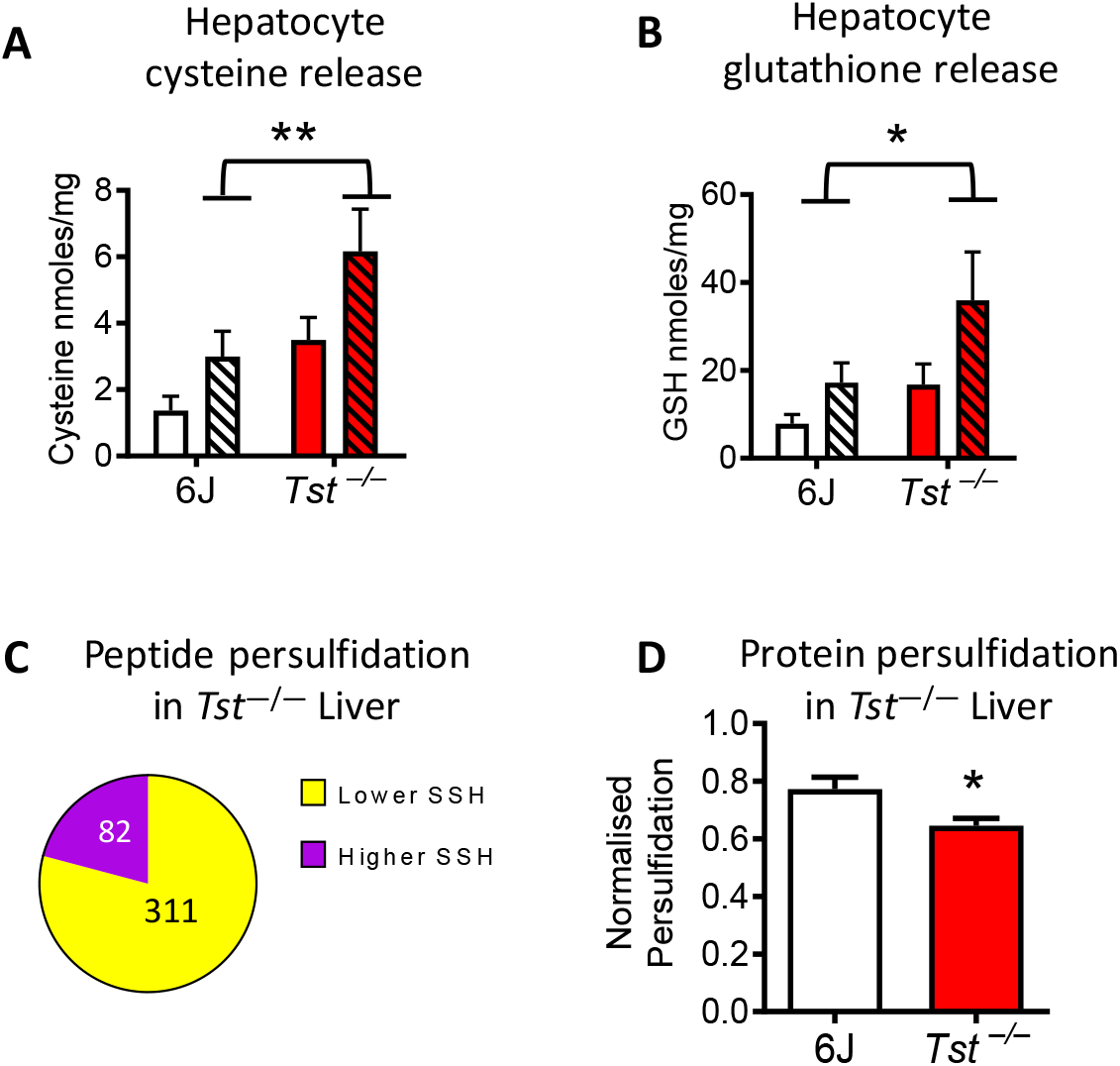
*Tst* deletion results in increased hepatic sulfur excretion and a reduction of protein persulfidation. (**A**) Cysteine concentrations (MBB-HPLC) in the media incubated with primary hepatocytes in the presence (hatched pattern) or absence (no pattern) of 1mM methionine, from C57Bl/6J (white bars, n = 4/treatment) and *Tst^−/−^* ^(^red bars, n = 4/treatment) mice. (**B**) Glutathione concentrations (MBB-HPLC) in the media incubated with primary hepatocytes in the presence (hatched pattern) or absence (no pattern) of 1mM methionine, from C57Bl/6J (white bars, n = 4/treatment) and *Tst^−/−^* ^(^red bars, n = 4/treatment) mice. (**C**) Pie chart depicting the proportion of liver peptides that are significantly higher (82 peptides, purple space) or lower (311 peptides, yellow space), in their persulfidation rate in the *Tst^−/−^* (n = 3) relative to C57Bl/6J (n = 3) mice. (**D**) Total DTT-released cysteine-persulfidated liver protein as measured by REVERT total protein stain following western blotting, normalised to the total input protein of the sample from *Tst^−/−^* (red bar, n = 4) and C57Bl/6J (white bar, n = 4) mice. Data with error bars are represented as mean ±SEM. Significance was calculated using 2-WAY ANOVA (**A**, **B**) or student’s t-test (**D**), * P < 0.05, ** P < 0.01. For (**A**) and (**B**) the 2-WAY ANOVA reveals a main effect of genotype, indicated by * or ** on the histogram. A significant effect of methionine was also found for both (**A**) and (**B**) not indicated on the histogram. For (**C**) peptides were selected as being significant at a P-diff of 0.95 or greater.

**Table 1.** *Tst* deletion results in altered sulfur metabolites in blood and liver. (**A**) Sulfide dibimane, thiosulfate-MBB, measured by fluorescence detection following HPLC, from whole blood taken from trunk blood of ND-fed C57Bl/6J (n = 4) and *Tst^−/−^* (n = 4) mice. (**B**) Sulfide dibimane, thiosulfate-MBB, and rGSH-MBB, measured by fluorescence detection following HPLC, from EDTA-Plasma of ND-fed C57Bl/6J (n = 4) and *Tst^−/−^* (n = 4) mice. (**C**) Thiosulfate-MBB corrected for creatinine from 24 hour urine samples, taken from ND-fed C57Bl6/J (n = 4) and *Tst^−/−^* (n = 5) mice. (**D**) Sulfide dibimane, and thiosulfate-MBB, from whole blood taken from the inferior vena cava downstream of the hepatic vein of ND-fed C57Bl/6J (n = 3) and *Tst^−/−^* (n = 3) mice. (**E**) Sulfide dibimane, thiosulfate-MBB, rGSH-MBB, and cysteine-MBB from whole liver (n=4/genotype) of ND-fed C57Bl/6J (n = 4) and *Tst^−/−^* (n = 4) mice. (**F**) Fluorescence from cultured hepatocytes following incubation with P3 (sulfide reactive probe) from ND-fed C57Bl/6J (n = 4) and *Tst^−/−^* (n = 4) mice. (**G**) Ratio of Mito N/MitoA from the liver of ND-fed C57Bl/6J (n = 5) and *Tst^−/−^* (n = 5) mice. Data are represented as mean ±SEM. Significance was calculated using unpaired two-tailed student’s t-test. * P < 0.05, ** P < 0.01, *** P < 0.001, **** P < 0.0001

|  | C57Bl/6J | *Tst*<sup>-/-</sup> | *Tst*<sup>-/-</sup>/6J<br>ratio | Significance |
| --- | --- | --- | --- | --- |
| **A Trunk Blood (Micromolar)** |  |  |  |  |
| MBB-S (Sulfide) | 2.28 +/- 0.43 | 22.18 +/- 0.85 | 9.73 | **** |
| MBB-SSO3 (Thiosulfate) | n.d. | 6.25 +/- 3.17 | n.c. | ns |
| **B Trunk Plasma (Micromolar)** |  |  |  |  |
| MBB-S (Sulfide) | 1.88 +/- 0.64 | 24.50 +/- 2.02 | 13.03 | **** |
| MBB-SSO3 (Thiosulfate) | 3.99 +/- 0.99 | 80.29 +/- 13.6 | 20.12 | ** |
| MBB-GSH (reduced Glutathione) | 48.0 +/- 1.15 | 86.25 +/- 6.27 | 1.80 | *** |
| **C Urine (Micromoles/creatinine/24hrs)** |  |  |  |  |
| MBB-SSO3 (Thiosulfate) | 4.99 +/- 2.6 | 2374 +/- 319 | 475.75 | **** |
| **D Inferior Vena Cava (Micromolar)** |  |  |  |  |
| MBB-S (Sulfide) | 1.22 +/- 0.20 | 3.58 +/- 0.87 | 2.93 | ns (0.08) |
| MBB-SSO3 (Thiosulfate) | 6.58 +/- 4.51 | 88.3 +/- 13.0 | 13.42 | * |
| **E Liver (μmoles/kg wet liver)** |  |  |  |  |
| MBB-S (Sulfide) | 13 +/- 1 | 17 +/- 3 | 1.31 | ns |
| MBB-SSO3 (Thiosulfate) | 4 +/- 1 | 15 +/- 7 | 3.75 | ns |
| DNFB-GSH (reduced Glutathione) | 6470 +/- 380 | 6850 +/- 30 | 1.04 | ns |
| DNFB-Cysteine (Cysteine) | 82 +/- 13 | 67 +/- 11 | 0.82 | ns |
| **F Sulfide P3 fluorescence (A510nm/protein)** |  |  |  |  |
| Hepatocyte | 7.22 +/- 1.00 | 7.89 +/- 0.80 | 1.09 | ns |
| **G Mitochondrial sulfide (MitoA)** |  |  |  |  |
| Liver | 0.78 +/- 0.16 | 1.14 +/- 0.45 | 1.46 | ns |

\* P < 0.05, \*\* P < 0.01, \*\*\* P < 0.001 \*\*\*\* P < 0.0001

**Table 2.** Tst deletion results in differential persulfidation rate of liver proteins. Significant GO terms represented by peptides with different persulfidation rates in the ND-fed *Tst^−/−^* liver relative to C57Bl/6J. ‘**Direction**’ indicates whether the persulfidation is decreased or increased in *Tst^−/−^* relative to C57Bl/6J. ‘**Genes**’ indicates the number of genes in the *Tst^−/−^* that represent the changes driving the GO term.

| GO-ID | Name | Direction<br>(*Tst*<sup>-/-</sup> vs 6J) | Genes |
| --- | --- | --- | --- |
| GO terms Identified by log fold change |  |  |  |
| 0050660 | FAD binding | Decreased | 12 |
| 0008168 | Methyltransferase activity | Decreased | 9 |
| 0016741 | Transferase activity, transferring one-carbon groups | Decreased | 9 |
| 0008565 | Protein transporter activity | Decreased | 8 |
| 0008238 | Exopeptidase activity | Decreased | 7 |
| 0005777 | Peroxisome | Decreased | 7 |
| 0042579 | Microbody | Decreased | 7 |
| 0003995 | Acyl-CoA dehydrogenase activity | Decreased | 6 |
| 0008483 | Transaminase activity | Decreased | 6 |
| 0016769 | Transferase activity, transferring nitrogenous groups | Decreased | 6 |
| 0008757 | S-adenosylmethionine-dependent methyltransferase activity | Decreased | 6 |
| 0016655 | Oxidoreductase activity, acting on NADH/NADPH, quinone | Decreased | 5 |
| 0004177 | Aminopeptidase activity | Decreased | 5 |
| 0000059 | Protein import into nucleus, docking | Decreased | 3 |
| 0005643 | Nuclear pore | Decreased | 3 |
| 0031965 | Nuclear membrane | Decreased | 3 |
| 0044453 | Nuclear membrane part | Decreased | 3 |
| 0046930 | Pore complex | Decreased | 3 |
| 0015629 | Actin cytoskeleton | Decreased | 3 |
| 0016652 | Oxidoreductase activity, NADH/NADPH, NAD/NADP acceptor | Decreased | 3 |
| 0050662 | Coenzyme binding | Increased | 5 |
| 0016651 | Oxidoreductase activity, NADH/NADPH, | Increased | 5 |
| 0003954 | NADH dehydrogenase activity | Increased | 4 |
| 0008137 | NADH dehydrogenase (ubiquinone) activity | Increased | 4 |
| 0050136 | NADH dehydrogenase (quinone) activity | Increased | 4 |
| 0006739 | NADP metabolism | Increased | 3 |
| 0006769 | Nicotinamide metabolism | Increased | 3 |
| 0006733 | Oxidoreduction coenzyme metabolism | Increased | 3 |

### The hepatic proteome of Tst^−/−^ mice reveals a distinct molecular signature of altered sulfur and mitochondrial nutrient metabolism

To gain molecular insight into the mechanisms underlying the apparently diabetogenic liver phenotype in ND-fed *Tst^−/−^* mice, we performed a comparative iTRAQ proteomic analysis of whole liver with ND-fed C57BL/6J mice. Kyoto Encyclopedia of Genes and Genomes (KEGG) analysis (at significance threshold P < 0.05) revealed 4 up–regulated pathways in liver of *Tst^−/−^* mice related to amino acid metabolism, including sulfur amino acids, and sulfur metabolism (Table 3A). GO analysis (at threshold P < 0.05) revealed 95 significantly up–regulated terms in liver of *Tst^−/−^* mice (Table S3A). Of the top 14 up–regulated terms, 7 referred to amino acid metabolism and 1 referred to the organellar term *‘mitochondrion’*. KEGG pathway analysis revealed 27 down–regulated pathways in the liver of *Tst*^−/−^ mice (Table 3B) including Phase 1 and 2 detoxification pathways (Cytochrome P450s, Glutathione and Glucuronidation) and *‘Lysosome’*, *‘Protein processing in the Endoplasmic reticulum’* organellar terms. 213 GO terms were significantly down–regulated in *Tst^−/−^* mice (Table S4B). Among the most significant down–regulated terms were phase 2 detoxification pathways such as *‘glutathione binding’* and *‘glutathione transferase activity’* and *‘endoplasmic reticulum’*. We validated a representative subset of protein changes (Figures S5A and S5D), noting broadly consistent direction of change in the key pathways. At the single protein level, the most robust change we observed was increased MPST protein in whole liver (Figure S5A and S5D) and mitochondrial sub-fractions (Figure S5B and S5D). This change was remarkable as mRNA levels for *Mpst* were lower in *Tst^−/−^* mice (Figure S5C), likely as a result of loss of proximal *Mpst* promoter function; *Mpst* is a paralog of *Tst*^47^ juxtaposed approximately 1kb from the *Tst* gene. Protein levels for other sulfide producing and disposal enzymes were comparable between genotypes (Table S4). Given the evidence for glucose and lipid metabolic changes in the *Tst^−/−^* mice, we undertook a further focussed comparison of canonical proteins in these pathways (Table S5). Four GO terms were enriched in liver from *Tst^−/−^* mice, all of which were down– regulated; *‘Lipid metabolic process’, ‘fatty acid beta-oxidation’, ‘Acyl-CoA dehydrogenase activity’ and ‘Acyl-CoA hydrolase activity’* (Table S5). Notably, supportive of largely unchanged hepatic insulin sensitivity, canonical insulin–regulated proteins^48, 49^ were expressed comparably between genotypes (Tables S6 and S7).

**Table 3.** Tst deletion results in differential hepatic protein abundance of KEGG pathways. (**A**) Significant KEGG pathway terms represented by proteins that are more abundant in the ND-fed *Tst^−/−^* liver compared with ND-fed C57Bl/6J. (**B**) Significant KEGG pathway terms represented by proteins that are less abundant in the ND-fed *Tst^−/−^* liver compared with ND-fed C57Bl/6J. ‘**Genes**’ indicates the number of genes in the *Tst^−/−^* that represent the changes driving the KEGG pathway.

| Entry | Name | Genes | Significance |
| --- | --- | --- | --- |
| **A. Increased in ND *Tst*<sup>-/-</sup> liver** |  |  |  |
| 00250 | Alanine, aspartate and glutamate metabolism | 6 | *** |
| 00260 | Glycine, serine and threonine metabolism | 5 | ** |
| 00270 | Cysteine and methionine metabolism | 4 | * |
| 04122 | Sulfur relay system | 2 | * |
| **B. Reduced in ND *Tst*<sup>-/-</sup> liver** |  |  |  |
| 00980 | Metabolism of xenobiotics by cytochrome P450 | 12 | **** |
| 00982 | Drug metabolism – cytochrome P450 | 12 | **** |
| 05204 | Chemical carcinogenesis | 12 | **** |
| 00480 | Glutathione metabolism | 8 | *** |
| 00040 | Pentose and glucuronate interconversions | 5 | ** |
| 04142 | Lysosome | 6 | ** |
| 04390 | Hippo signalling pathway | 4 | ** |
| 00500 | Starch and sucrose metabolism | 5 | ** |
| 05215 | Prostate cancer | 3 | ** |
| 04024 | cAMP signalling pathway | 4 | * |
| 04141 | Protein processing in ER | 9 | * |
| 05211 | Renal cell carcinoma | 3 | * |
| 00830 | Retinol metabolism | 6 | * |
| 00053 | Ascorbate and aldarate metabolism | 4 | * |
| 00860 | Porphyrin and chlorophyll metabolism | 4 | * |
| 04722 | Neurotrophin signalling pathway | 3 | * |
| 04670 | Leukocyte transendothelial migration | 4 | * |
| 04010 | MAPK signalling pathway | 4 | * |
| 04720 | Long-term potentiation | 2 | * |
| 04914 | Progesterone-mediated oocyte maturation | 2 | * |
| 04062 | Chemokine signalling pathway | 3 | * |
| 04110 | Cell cycle | 3 | * |
| 04015 | Rap1 signaling pathway | 4 | * |
| 00983 | Drug metabolism - other enzymes | 5 | * |
| 04918 | Thyroid hormone synthesis | 3 | * |
| 04612 | Antigen processing and presentation | 3 | * |
| 05203 | Viral carcinogenesis | 5 | * |
| * P < 0.05, ** P < 0.01, *** P < 0.001 **** P < 0.0001 |  |  |  |

### Hepatic protein expression in Tst^−/−^ mice is consistent with lower NRF2 activation

We performed a transcription factor binding site (TFBS) enrichment analysis in the promoters of proteins that are up–regulated in the liver of *Tst^−/−^* mice to look for potential hub transcriptional drivers of the proteome profile (Figure S6A). This revealed a statistically significant under-representation of TFBS for the sulfide–responsive^50,51^ NRF2 transcription factor (Figure S6A). To explore whether there was evidence for reduced NRF2 activation in the *Tst^−/−^* liver, we looked at the relative expression of 47 NRF2–regulated proteins in the proteome of ND-fed *Tst^−/−^* and ND-fed C57Bl/6J mice. While 37 of these target proteins were similar in expression between the genotypes, 10 NRF2 target proteins were lower in the liver of *Tst^−/−^* mice (Figure S6B) and none of the 47 NRF2 target proteins were increased in *Tst^−/−^* liver.

### Proteome of TST deficiency versus HFD response in C57BL/6J mice reveals distinct regulation of lipid metabolism, sulfide metabolism and detoxification pathways

We examined mechanistic commonalities between the diabetogenic hepatic phenotype of *Tst^−/−^* mice and that induced by the diabetogenic HFD–feeding regimen in C57BL/6J mice. Note, ND-fed *Tst^−/−^* mice were in a pre-existing diabetogenic state (Figure 1) that does not further worsen with HFD (Figure 1H-I, Table S1), suggesting gross phenotypic convergence of the two genotypes after HFD. We compared the identity and direction of change of the 188 proteins differentially expressed in ND-fed *Tst^−/−^* mice (versus ND-fed C57BL/6J mice; Figure 3A) to proteins that were differentially expressed in response to HFD in C57BL/6J mice (432 proteins; Figure 3A). There was a striking 67% overlap in individual proteins (126) in this comparison (Figure 3A). When we analysed these two protein signatures for directionally shared pathways, one up-regulated KEGG pathway *‘Glycine, serine and threonine metabolism’* (Table S8A) and 12 down-regulated KEGG pathways, including *‘drug metabolism’* and *‘endoplasmic reticulum’* (Table S8B) were common to the liver of the ND-fed *Tst^−/−^* and HFD-fed C57BL/6J mice. Consistent with a pre-existing HFD–like proteome, the dynamic response to HFD in the liver of *Tst^−/−^* mice was muted, relative to that observed in C57BL6J mice (106 proteins, a 4–fold lower response; Figure 3B). Focussing on the sulfide pathway, MPST and SUOX were increased by HFD in 6J and *Tst^−/−^* mice (Table S9). The HFD–induced increase in MPST was less pronounced in the liver of *Tst^−/−^* mice, likely a reflection that it is already elevated in ND–fed *Tst^−/−^* mice. We then considered contrasting, rather than congruent, proteomic responses arising from TST deficiency versus HFD responses in C57BL/6J mice to illuminate potential novel pathways underlying the otherwise functionally similar diabetogenic hepatic *Tst^−/−^* phenotype. 5 KEGG pathways (Table S10A) and 4 GO terms (Table S10B) were regulated in the opposite direction in this comparison. Strikingly, the GO terms were all related to lipid metabolism, which were up–regulated in the HFD response but down– regulated with TST deficiency (Table S10A-B). An organelle–focussed protein analysis showed shared up-regulation of mitochondrial and endoplasmic reticulum pathways between TST deficiency (Figure 3C upper row) and C57BL/6J HFD-responses (Figure 3C lower row), but a striking discordance in peroxisomal protein pathways (up-regulated by HFD, down-regulated TST deficiency) and nuclear proteins (down-regulated by HFD, up-regulated TST deficiency; Figure 3C).

**Figure 3.**
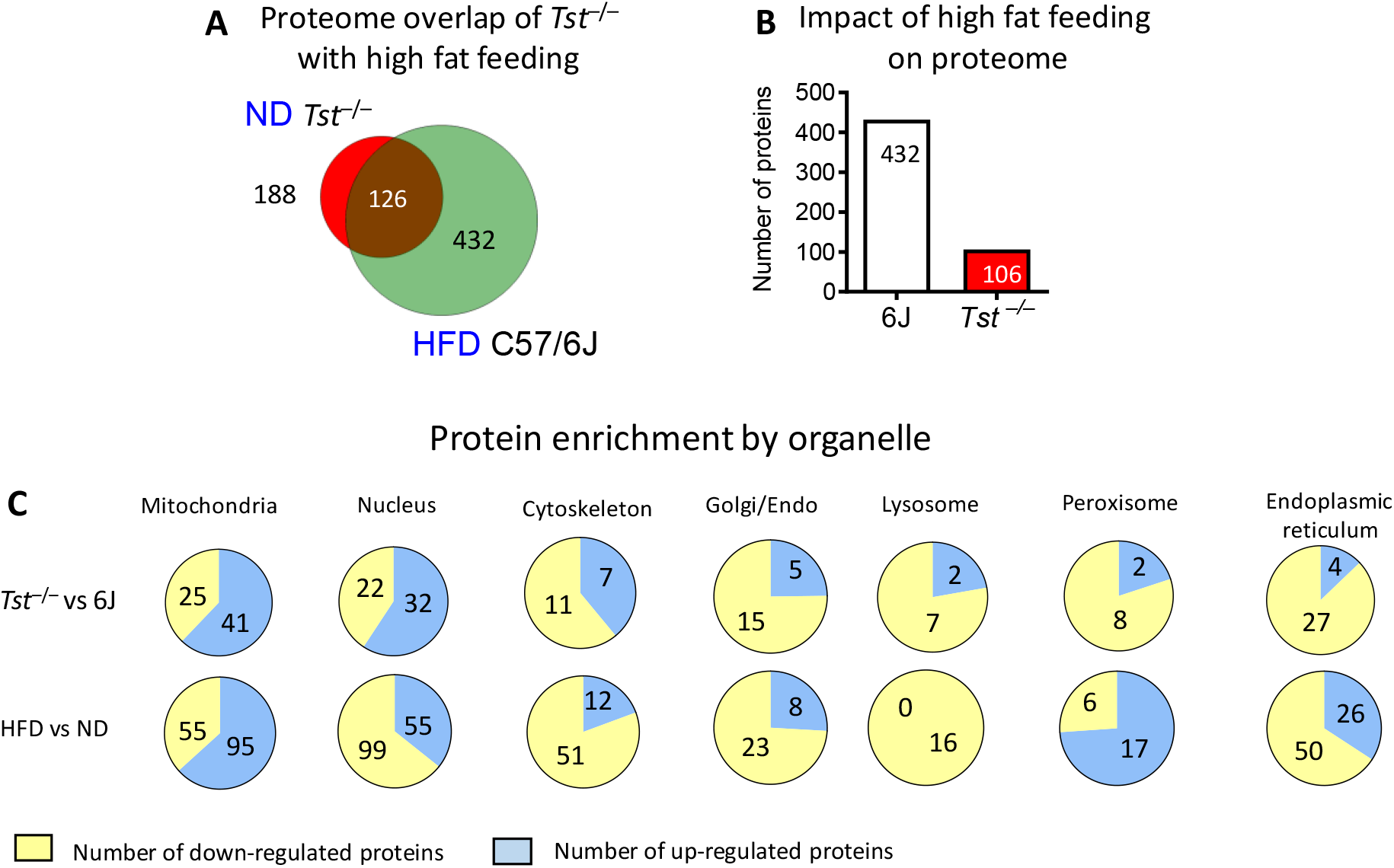
*Tst* deletion engenders a high fat feeding–like hepatic proteome with a distinct organellar signature. (**A**) Venn diagram representing the number of proteins significantly different (at P < 0.01) between normal diet-fed *Tst^−/−^* and C57Bl/6J (Red circle), and the number of regulated proteins between high fat fed and normal diet-fed C57Bl/6J (Green circle). The overlap (brown) represents those proteins regulated in the same direction by both comparisons (n= 4/genotype). (**B**) Number of proteins significantly different (at p < 0.01) between 58% high fat and normal diet in either C57Bl/6J (white bar), or *Tst^−/−^* mice (Red bar), (n= 4/genotype). (**C**) Pie charts depicting the proportion of individual liver proteins that are upregulated (Blue space) compared to downregulated (yellow space) after GO term categorisation according to subcellular location. Upper row; normal diet-fed *Tst^−/−^* relative to normal diet-fed C57Bl/6J. Lower row, high fat-fed C57Bl/6J relative to normal diet-fed C57Bl/6J.

### The Tst^−/−^ liver proteome and persulfidome converge on transamination and lipid oxidation pathways

To assess whether conservation of changes at protein and post-translational modification levels can illuminate key regulatory hubs driving the hepatic phenotype we ran a congruence analysis of the proteome and persulfidome. We found that GO terms referring to *amino acid, lipid metabolism* and *peroxisome* were significantly regulated at both the level of protein abundance and persulfidation status (Table 4).

**Table 4.** The *Tst^−/−^* liver proteome and persulfidome converges on lipid oxidation pathways. GO terms that are significantly regulated following comparison of ND-fed *Tst^−/−^* relative to C57Bl/6J, at *both* the level of cysteine persulfidation and protein abundance.

| Table 4. GO Terms: Congruence of peptide abundance and persulfidation |  |  |  |
| --- | --- | --- | --- |
| GO-ID | Name | Persulfidation<br>(*Tst*<sup>-/-</sup> vs 6J) | Abundance<br>(*Tst*<sup>-/-</sup> vs 6J) |
| 0008483 | Transaminase activity | Decreased | Increased |
| 0016769 | Transferase activity, transferring nitrogenous | Decreased | Increased |
| 0003995 | Acyl-CoA dehydrogenase activity | Decreased | Decreased |
| 0005777 | Peroxisome | Decreased | Decreased |
| 0042579 | Microbody | Decreased | Decreased |

### Tst^−/−^ hepatocytes exhibit elevated mitochondrial respiration and a defect in medium–chain fatty acid oxidation

Enhanced respiratory sulfide disposal was found from *Tst^−/−^* hepatocytes, and enrichment of mitochondrial proteins was suggested from the liver proteome of the *Tst^−/−^* mice (Table S4). We therefore sought to determine if TST deficiency affected respiratory function and substrate utilisation of the hepatocyte. Analysis of electron micrographs prepared from the liver of ND-fed *Tst^−/−^* mice and C57BL/6J controls showed morphologically normal mitochondria (Figure 4A). Basal respiration, comprising ATP–linked and leak respiration, was significantly higher in hepatocytes from *Tst^−/−^* mice (Figure 4B–4D). Maximal hepatocyte respiratory capacity and non–respiratory oxygen consumption was similar between genotypes (Figures S7A and S7B). In line with phenotypic convergence following HFD, hepatocyte respiration was comparable between genotypes from HFD-fed mice (Figures S7C-S7H). A unique feature of the liver from *Tst^−/−^* mice that contrasted with HFD response in C57BL/6J mice was a decrease in proteins and persulfidation levels of proteins in lipid oxidation pathways. We therefore investigated hepatocyte respiration of lipids. Using a low pyruvate (100 μM) medium to reveal respiratory dependency on other substrates, we showed that CPT1A–mediated mitochondrial oxidation of endogenous long chain fatty acids (LCFA; etomoxir-inhibited) was similar between genotypes (Figure 4E). Next, we by–passed CPT1A–mediated LCFA transfer, and revealed a marked deficit in respiration stimulated by the medium chain fatty acid octanoate in hepatocytes from *Tst^−/−^* mice (Figure 4F). A similar experiment adding back pyruvate revealed comparable stimulation of respiration between genotypes (Figure S6I). In amino acid free media, combined glutamine–, aspartate– and alanine–stimulated hepatocyte respiration was comparable between genotypes (Figure S6J).

**Figure 4.**
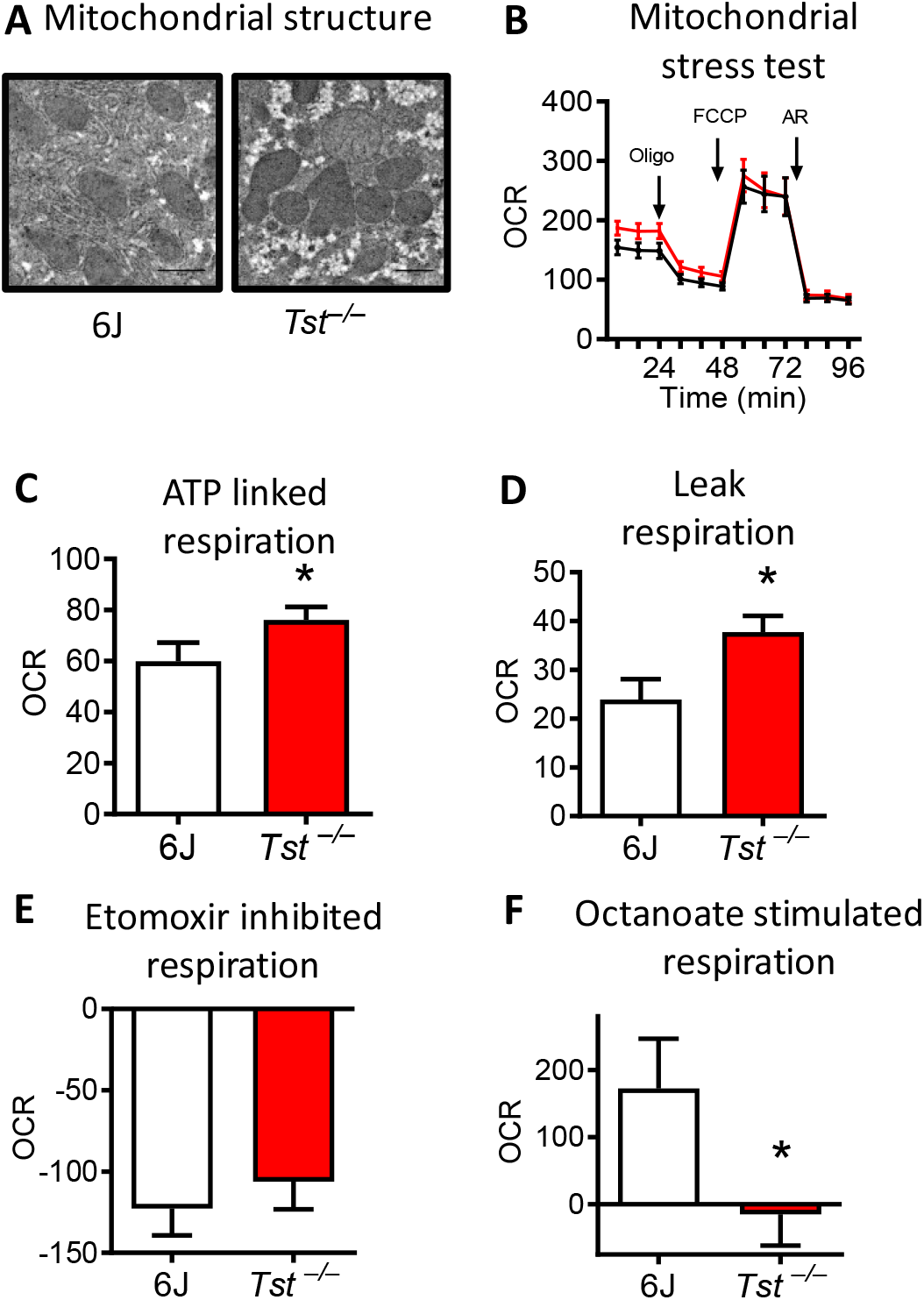
*Tst* deletion results in increased hepatocyte respiration but impaired medium–chain fat respiration. (**A**) Electron microscope images of liver, visualising mitochondria from normal diet-fed C57Bl/6J (n = 4) or *Tst^−/−^* (n = 4) mice. (**B**) Seahorse trace representing the mean oxygen consumption rate (OCR), normalised to protein, by hepatocytes from normal diet-fed C57Bl/6J (n = 6) or *Tst^−/−^* (n = 6) mice during a mitochondrial stress test. (**C**) Respiratory OCR linked to ATP production (oligomycin sensitive) by hepatocytes from normal diet-fed C57Bl/6J (n = 6) or *Tst^−/−^* (n = 6) mice, calculated from Figure 4B. (**D**) Respiratory OCR relating to proton leak (oligomycin insensitive) by hepatocytes from normal diet-fed C57Bl/6J (n = 6) or *Tst^−/−^* (n = 6) mice, calculated from Figure 4B. (**E**) Reduction of maximal uncoupled respiration following inhibition of long chain fatty acid mitochondrial import using etomoxir (8 μM), from normal diet-fed C57Bl/6J (n = 4) or *Tst^−/−^* (n = 4) mice. (**F**) Stimulation of maximal uncoupled respiration following addition of medium chain fatty acid octanoate (250 μM), from normal diet-fed C57Bl/6J (n = 4) or *Tst^−/−^* (n = 4) mice. Data are represented as mean ±SEM. Significance was calculated using an unpaired two tailed, student’s t-test (**C**, **D**, **E**, **F**), * P < 0.05.

## Discussion

Elevated TST expression in adipose tissue was recently identified as a genetic mechanism driving metabolically protective leanness in mice^35^. Conversely, mice with genetic deletion of *Tst^−/−^* (*Tst*^−/−^ mice) exhibited impaired glucose tolerance relative to C57BL/6N controls^35^. A key observation from the *Tst^−/−^* mouse was qualitative evidence for elevated blood sulfide levels^35^, which implicated the mitochondrial sulfurtransferase as playing an important role in the sulfide oxidation pathway (Figure S8) for the first time *in vivo*. However, the mechanistic link between *Tst^−/−^* deficiency and impaired glucose tolerance remained unclear. The *Tst* mice had a subtle adipose tissue phenotype, with comparable fat accumulation upon high fat feeding, suggesting alternate organs as a potential source of impaired glucose homeostasis. Moreover, TST is most highly abundant in the liver, consistent with its role in detoxification of cyanide and related molecules. Indeed, hepatic TST mRNA levels are ~8-fold higher than adipose tissue, with the difference in enzyme activity likely substantially exceeding this, per gram of tissue. Based upon the reported effects of sulfide to increase hepatic glucose production^20,21^ and modulate lipid metabolism^22,23^, we hypothesised that TST deficiency predominantly drives a diabetogenic state through increased hepatic sulfide exposure. Consistent with this, we found evidence supportive of increased gluconeogenesis, steatosis, elevated plasma VLDL triglycerides and, quantitatively, a 10-fold higher circulating blood sulfide level in *Tst^−/−^* mice. Unexpectedly, and in spite of the markedly increased circulating sulfide levels, steady-state sulfide level was normal in the liver of *Tst^−/−^* mice. Moreover, we found evidence for multiple mechanisms for increased, not decreased, hepatic sulfide disposal, reduced downstream sulfide signalling and novel molecular explanations for the apparently diabetogenic phenotype. Our data are consistent with the liver being a major site for sulfide disposal rather than production^32^. Moreover, our data suggest that the liver of *Tst^−/−^* mice has over-compensated in its response to maximise hepatic sulfide removal, thus engendering detrimental metabolic consequences indirectly. This response is achieved through a combination of distinct compartmentalised cellular responses including increased respiratory sulfide disposal and export of cysteine and GSH and is further evidenced by the reduction of global hepatic protein persulfidation and depletion of sulfide-responsive^51^ NRF2 target proteins. Additionally, up-regulation of translation and recruitment of MPST to the mitochondria of *Tst^−/−^* mice is observed. This response, in face of reduced transcription of *Mpst* due to *Mpst* promoter disruption by the *Tst* null allele, suggests a powerful post-transcriptional cellular sulfide sensing mechanism. Moreover, if MPST is compensating for TST–mediated sulfide disposal in this context, this suggests a subversion of normal MPST function away from sulfide production^52,53,54,55^. Intriguingly, TST expression levels were elevated in the liver of *Mpst^−/−^* mice, providing further support for an exquisite reciprocal compensatory mechanism between these two enzymes^56^.

The unexpected finding of normal hepatic sulfide levels in the *Tst^−/−^* mice led us to discover that the metabolic phenotype we observed was driven by the very mechanisms invoked to maintain sulfide within a normal range rather than sulfide excess *per se*. Several observations were consistent with this. For example, the major amino acid pathways increased in the liver proteome of *Tst^−/−^* mice were transaminases involved in metabolism of GSH that support increased export of sulfur equivalents as GSH (and cysteine). These same transaminases support gluconeogenesis by redirecting Krebs cycle intermediates^57–59^. Re–programming of amino acid metabolism for sulfide disposal, with knock–on effects to drive hepatic glucose production are suggested, rather than any change to amino acid–linked mitochondrial respiration in hepatocytes. This is supported by the shift in pyruvate metabolism towards aspartate. In addition, glutathione-S-transferases (GST) that inhibit gluconeogenesis^60^ were lower in the liver of *Tst^−/−^* mice. Further, NRF2 activation, which represses gluconeogenesis^61^ appears lower in liver of *Tst^−/−^* mice. The involvement of NRF2 in the *Tst^−/−^* liver phenotype is further supported by the phenotype of *Nrf2^−/−^* mice that also exhibited steatohepatitis, in the absence of insulin resistance^62^. However, we note that NRF2 signalling can be complex and dependent upon dietary context; *Nrf2^−/−^* mice showed improved glucose tolerance after a high–fat diet^63^ suggesting any contribution of a NRF2 signalling deficit in the liver of the *Tst^−/−^* mice changes upon high fat feeding. Beyond altered pyruvate flux, we also showed that the hepatocytes of *Tst^−/−^* mice exhibited defective lipid metabolism. Specifically, medium–chain fatty acid (MCFA) oxidation was impaired, associated with selective reduction of protein levels and persulfidation levels of lipid catabolic enzymes. This represents a novel mechanism linking altered sulfide metabolism to lipid oxidation, hepatic lipid accumulation and dyslipidaemia. Consistent with impaired MCFA oxidation defects as a driver, at least in part, of the phenotype of *Tst^−/−^* mice, steatosis is observed in medium–chain acyl-CoA dehydrogenase (*Mcad*)^−/−^ mice^64^ and dyslipidemia is found in MCADD deficient humans^65^. The data we present adds to a growing understanding of the link between sulfide regulating genes and nutrient metabolism that has hitherto focussed on the enzymes of sulfide production. Specifically, we provide support for the importance of the sulfide oxidising pathway as a regulator of cellular sulfide exposure. Unexpectedly, the data reveal novel cellular mechanisms that are engaged to homeostatically regulate sulfide disposal and that can impact upon cell energetics and nutrient metabolism.

Our findings may have implications for potentially unexpected side effects of sulfide donor therapeutics. In normal mice, *in vivo* sulfide administration for 4 weeks post–HFD partially reversed hepatic lipid accumulation invoked by chronic (16 weeks) HFD^22^. No evidence was provided for whether sulfide disposal mechanisms were altered^22^. This efficacious sub-chronic sulfide administration regimen contrasts with our genetic model of chronic sulfide elevation as a driver of dysregulated metabolism and NAFLD. Clearly, the normal mice in the Na2S administration studies had a fully functional SOP, suggesting the presence of TST is required to achieve the beneficial metabolic effects of Na2S administration. This is also consistent with the apparently low sulfide signalling status (evidenced by lower persulfidation, NRF2 target protein abundance) in the liver of the *Tst^−/−^* mice. The benefits of elevated sulfide cannot be realised, perhaps because the major mediator of those effects is missing and the alternate mechanisms invoked do not fully compensate (e.g. MPST), or actively drive aberrant nutrient metabolism. Comparable studies of glucose and lipid metabolism after manipulation of other sulfide regulating genes are limited. However in a contrasting model of reduced sulfide production (*Cth^−/−^* mice), plasma triglycerides were lowered^23^, consistent and opposite to what we observed with the *Tst^−/−^* mice. The hepatic sulfide disposal status of the *Cth*^−/−^ mouse model is unknown, but our findings predict a suppression of the SOP to spare the limited endogenous sulfide produced. Intriguingly they also predict a knock-on effect on nutrient homeostasis due to reduced metabolic demand of the TST/SOP axis. A more direct model informing on the effects of impairment of the sulfide disposal pathway is deficiency of the key mitochondrial SOP enzyme ETHE1. *Ethe1^−/−^* mice suffer fatal sulfide toxicity^30^ and therefore comparable metabolic studies are lacking. However, one notable observation is that *Ethe1^−/−^* mice have an apparently 10-fold higher liver sulfide exposure than control mice^30^, in contrast to the normalised hepatic sulfide levels of *Tst^−/−^* mice. Circulating sulfide levels were not reported for comparison, but the presumably relatively lower systemic sulfide levels of *Tst^−/−^* mice appear to have permitted an effective homeostatic sulfide disposal response in the liver to avoid toxicity, albeit with a metabolic cost. Consequently, the liver of *Tst^−/−^* mice has a distinct functional and proteomic profile to that of the *Ethe1^−/−^* mice. For example, in the liver of *Tst^−/−^* and *Ethe1^−/−^* mice^66^, proteins of the glutathione–S–transferase Mu type (GSTM) and peroxiredoxin (PRDX) families were altered, but sometimes in the opposite direction or with alteration of distinct protein sub-classes. A notable difference is also observed in amino acid metabolism. The liver of *Ethe1^−/−^* mice exhibited increased expression of enzymes of branched chain amino acid metabolism^66^, distinct from the predominantly glutathione–related amino acid pathways that are increased in liver of *Tst^−/−^* mice. Beyond sulfide, TST may also have distinct cellular roles that affect metabolism such as mitoribosomal synthesis, ROS attenuation and modulation of mitochondrial iron-sulfur clusters^67–71^.

Given the pro-diabetogenic liver phenotype in *Tst^−/−^* mice, its was surprising that insulin signalling in the liver appeared normal, and peripheral insulin sensitivity was, if anything, increased. There are precedents for increased hepatic glucose production independent of insulin resistance, as found in the *Nrf2^−/−^* mice^62^ and as driven by the transcription factor ChREBP^72,73^. There is also evidence to support insulin–sensitising effects of sulfide administration *in vivo* in mice and rats^74–76^, consistent with a sulfide–mediated insulin–sensitisation of non-hepatic tissues in *Tst^−/−^* mice. Higher circulating GSH in *Tst^−/−^* mice may also promote peripheral insulin–sensitisation^77,78^. Clearly, the net balance of glucose production from the liver and its peripheral disposal remains abnormal in *Tst^−/−^* mice. Indeed, the baseline metabolic phenotype of *Tst^−/−^* mice resembles in many ways that of a normal mouse fed a HFD. Thus, we expected to gain insight into conserved pro-diabetogenic mechanisms from the overlap between the liver proteome of *Tst^−/−^* mice and that of HFD-fed C57BL/6J mice. However, upon closer inspection this was not entirely the case, with some lipid metabolism and peroxisomal proteins changing in the opposite direction to those observed in HFD–fed C57BL/6J mice. Unlike a HFD state, which is associated with dominant hepatic insulin resistance, the increased hepatic glucose production in ND-fed *Tst^−/−^* mice occurs despite of normal hepatic insulin sensitivty. The significant changes in persulfidation of transaminase and gluconeogenesis proteins in the liver of *Tst^−/−^* mice suggest a coordinated cross–talk across metabolic pathways underlies this atypical metabolic phenotype.

Sulfide donor therapeutics were proposed as a clinical strategy for improving cardiovascular health^27,79,80^. Elevated endogenous sulfide was also implicated in the beneficial metabolic effects of caloric restriction^11,81–85^. Our results suggest that chronic sulfide elevation may have unintended detrimental consequences, driving liver glucose production and fat accumulation to undesirable levels. This caveat may be fortunately limited to cases where SOP proteins are compromised through rare genetic effects – such as TST variants^86,87^. More broadly, a number of drugs or supplements are known to increase cyanide, which may dominantly inhibit TST activity and result in secondary sulfide overexposure. These include nitroprusside^88^ and amygdalin^89,90^. Indeed the TST metabolite thiosulfate is commonly co-administered with nitroprusside to prevent cyanide toxicity^91^. Furthermore, dietary and environmental exposure to cyanogenic compounds^92^, e.g. smoking^93^ or cyanogenic diets^94^ may interfere with normal TST function and could lead to increased sensitivity to sulfide therapeutics. In contrast, we have shown that administration of the TST substrate thiosulfate can ameliorate diabetes^35^ further underlining the potential utility of targeting the SOP in metabolic disease. As with all therapeutic strategies, a careful cost-benefit analysis is required. A comparable case of relevance are the statins, one of the most potent and widely used drugs to prevent atherosclerosis, which also carry a higher risk for diabetes^95^. The full impact of TST manipulation on opposing metabolic pathways requires further study. Our current study sheds light on the underlying hepatic mechanisms invoked for sulfide disposal that are relevant to current sulfide–donor strategies and may inform on routes to reduce their potential metabolic side-effects.

## Methods

The methods for all data and work presented in this manuscript is provided as supplementary information.

## Author Contributions

N.M.M. and R.N.C conceived experiments. R.N.C., M.T.G.G, M. B.-L., A. M.-C., M.L., V.V., B.E., T. LeB., M.B., S.H., S.D., N.H., C. Mc F., A.T., N. F., T.G., performed experiments. R.N.C., M. B.-L., P.F., V.V., T. LeB., N.H., T.S., F.B., T.G., R.C.H., B.S., G.G., A.J.F., C.S., R.B. and N.M.M. analysed and interpreted data and commented on the manuscript. K.H.A., S.S. generated reagents. C.McM. and R.C.H. generated reagents. R.N.C. and N.M.M. wrote the manuscript.

## Acknowledgements

This work was funded by a Wellcome Trust New Investigator Award to NMM (100981/Z/13/Z) and Diabetes UK grant (17/0005697). RCH was funded by a Medical Research Council Discovery Award (MC-PC-15076). We would like to thank Professor Ken Olsen and Eric DeLeon (Notre Dame) for advice establishing sulfide measurement by probe.

