## Supplementary Information for "The hepatic compensatory response to elevated systemic sulfide promotes diabetes"

### Supplementary Tables

**Table S1. Parameters during the euglycemic hyperinsulinemic clamp**

**(A) Parameters during the basal (pre clamp) experiment (60-90 minutes post tracer)**

| Parameter | 6J chow | *Tst*$^{-/-}$ chow | 6J HFD | *Tst*$^{-/-}$ HFD | Genotype | Diet |
|---|---|---|---|---|---|---|
| Fasted Glucose 60 min (mg/dl) | 116.35± 14.93 | 135.72± 7.22 | 146.50± 3.89 | 167.13± 9.68 | * | ** |
| Glycolysis (mg/kg/min) | 11.63 ± 1.60 | 11.12 ± 0.62 | 12.89 ± 0.57 | 12.61 ± 0.62 | ns | ns (0.09) |
| Glycogen synthesis (mg/kg/min) | 21.48 ± 2.06 | 19.02 ± 2.04 | 15.08 ± 2.76 | 16.28 ± 2.50 | ns | **** |

**(B) Measurements and parameters during the clamp experiment (160-210 minutes post tracer)**

| Parameter | 6J chow | *Tst*$^{-/-}$ chow | Genotype (chow) | 6J HFD | *Tst*$^{-/-}$ HFD | Genotype (HFD) |
|---|---|---|---|---|---|---|
| Glucose 160 min (mg/dl) | 108.0 ± 5.0 | 121.0 ± 10.1 | ns | 120.1± 3.7 | 125.6 ± 10.3 | ns |
| Glucose 170 min (mg/dl) | 120.3 ± 6.4 | 129.8 ± 5.8 | ns | 118.1 ± 4.1 | 143.9 ± 14.0 | ns (0.08) |
| Glucose 180 min (mg/dl) | 115.0 ± 5.5 | 125.5 ± 2.5 | ns (0.08) | 125.9 ± 2.9 | 126.7 ± 6.8 | ns |
| Glucose 190 min (mg/dl) | 124.0 ± 1.0 | 131.0 ± 4.6 | ns | 121.5 ± 4.7 | 116.1 ± 3.7 | ns |
| Glucose 200 min (mg/dl) | 121.0 ± 5.0 | 125.3 ± 5.8 | ns | 121.9 ± 3.0 | 115.7 ± 5.3 | ns |
| Glucose 210 min (mg/dl) | 113.0 ± 7.8 | 123.5 ± 4.6 | ns | 119.4 ± 4.0 | 116.9 ± 2.7 | ns |
| Glucose IR 160-210 min (mg/kg/min) | 84.9 ± 4.2 | 85.0 ± 2.6 | ns | 70.26 ± 4.83 | 69.95 ± 4.96 | ns |
| Glucose IR 160 (mg/kg/min) | 82.3 ± 5.1 | 87.1 ± 3.2 | ns | 68.8 ± 4.2 | 68.9 ± 5.1 | ns |
| Glucose IR 170 (mg/kg/min) | 85.2 ± 4.9 | 89.0 ± 3.0 | ns | 70.0 ± 5.1 | 74.6 ± 4.4 | ns |
| Glucose IR 180 (mg/kg/min) | 84.7 ± 4.4 | 83.5 ± 4.2 | ns | 70.6 ± 5.1 | 70.7 ± 6.2 | ns |
| Glucose IR 190 (mg/kg/min) | 85.2 ± 4.0 | 86.1 ± 2.1 | ns | 69.9 ± 4.8 | 69.1 ± 5.7 | ns |
| Glucose IR 200 (mg/kg/min) | 85.2 ± 4.0 | 84.8 ± 2.4 | ns | 70.4 ± 4.7 | 71.2 ± 4.5 | ns |
| Glucose IR 210 (mg/kg/min) | 85.2 ± 4.0 | 84.5 ± 2.4 | ns | 70.4 ± 4.7 | 71.2 ± 4.5 | ns |
| Turnover (mg/kg/min) | 85.97 ± 3.52 | 94.50 ± 3.87 | ns | 73.61 ± 5.07 | 63.08 ± 7.10 | ns |
| Hepatic Glucose Prod. (mg/kg/min) | 1.10 ± 5.31 | 9.92 ± 9.15 | ns | 3.326 ± 4.03 | -6.83 ± 7.93 | ns |
| Glycolysis (mg/kg/min) | 45.90 ± 2.218 | 48.37 ± 2.05 | ns | 42.85 ± 1.48 | 36.42 ± 4.62 | ns (0.19) |
| Glycogen synthesis (mg/kg/min) | 40.06 ± 4.44 | 46.12 ± 2.68 | ns | 30.76 ± 5.36 | 26.66 ± 4.49 | ns |
| Integral Glucose (dpm.min/mg) | 3.6e$^{7}$ ± 1.4e$^{6}$ | 2.95e$^{7}$ ± 1.2e$^{6}$ | * | 1.88e$^{7}$ ± 1.4e$^{6}$ | 1.7e$^{7}$ ± 1.6e$^{6}$ | ns |
| IWAT glucose utilization (ng/mg.min) | 14.21 ± 4.05 | 18.62 ± 2.04 | ns | 4.53 ± 1.07 | 6.64 ± 1.00 | ns |
| EWAT glucose utilization (ng/mg.min) | 7.523 ± 4.39 | 7.21 ± 2.18 | ns | 2.94 ± 0.56 | 3.64 ± 0.39 | ns |
| VL glucose utilization (ng/mg.min) | 33.77 ± 2.98 | 38.66 ± 1.70 | ns (0.17) | 49.65 ± 9.19 | 47.42 ± 9.19 | ns |
| EDL glucose utilization (ng/mg.min) | 35.79 ± 11.09 | 40.86 ± 10.25 | ns | 68.79 ± 8.65 | 61.29 ± 11.80 | ns |
| Soleus glucose utilization (ng/mg.min) | 99.85 ± 12.38 | 123.60 ± 13.90 | ns (0.13) | 220.8 ± 24.45 | 198.5 ± 32.92 | ns |
| Tibialis glucose utilization (ng/mg.min) | 47.25 ± 7.22 | 53.67 ± 4.40 | ns | 74.39 ± 6.46 | 80.16 ± 6.09 | ns |
| Heart glucose utilization (ng/mg.min) | 161.40 ± 7.65 | 198.00 ± 14.09 | ns (0.13) | 226.6 ± 51.01 | 262.5 ± 23.29 | ns |
| Liver glucose utilization (ng/mg.min) | 3.53 ± 0.56 | 3.57 ± 0.63 | ns | 3.378 ± 0.39 | 3.58 ± 0.46 | ns |
| End Clamp Insulin (µU/ml) | 133.6 ± 7.03 | 126.4 ± 5.06 | ns | 147.2 ± 9.4 | 124.4 ± 10.9 | ns |

*** P < 0.05, ** P < 0.01, *** P < 0.001, **** P < 0.0001**

**Table S1. Parameters during the euglycemic hyperinsulinemic clamp.** Metabolic parameters measured during continuous trace infusion but prior to clamp (**A**) and during maintenance of euglycemia and hyperinsulinemia (**B**) from C57Bl/6J (chow-fed, n = 3, hfd-fed, n = 8) and *Tst*$^{-/-}$ (chow-fed, n = 6, hfd-fed, n = 7) mice. Data are represented as mean ±SEM. Significance for the basal experiment (**A**) was calculated using a 2-WAY ANOVA for *genotype* and *diet*. Significance for the clamp (**B**) was calculated for each diet separately using T-tests for *genotype*. * P < 0.05, ** P < 0.01, *** P < 0.001, **** P < 0.0001

| Table S2. Hydrogen sulfide disposal by hepatocytes and mitochondria (Amperometry) | | | |
|---|---|---|---|
| ***nmoles/min/mg protein*** | **C57Bl/6J** | $Tst^{-/-}$ | **Significance** |
| Hepatocytes | 3.88 +/- 0.095 | 4.15 +/- 0.345 | **ns** |
| Hepatocytes (Respiratory) | 1.97 +/- 0.176 | 2.91 +/- 0.288 | * |
| Hepatocytes (Non-respiratory) | 1.91 +/- 0.181 | 1.24 +/- 0.117 | * |
| Liver Mitochondria | 0.65 +/- 0.095 | 1.23 +/- 0.129 | * |
| Liver Mitochondria (Respiratory) | 0.26 +/- 0.060 | 0.50 +/- 0.080 | * |
| *** P < 0.05** | | | |

**Table S2. *Tst* deletion results in increased respiratory $H_2S$ disposal by hepatocytes**. $H_2S$ disposal rates (measured by gas selective amperometry following addition of 10 µM $Na_2S$) of hepatocytes (n = 6/genotype), or isolated liver mitochondria (n = 7/genotype) of ND-fed C57Bl/6J and $Tst^{-/-}$mice. Rates of $H_2S$ disposal were measured with and without respiratory inhibition following addition of Antimycin (2µM). Antimycin insensitive disposal rates are referred to as non-respiratory. The Antimycin sensitive disposal rates are referred to as respiratory. Data are represented as mean ±SEM. Significance was calculated using paired two-tailed student's t-test.* P < 0.05.

**Table S3 Extended list of significant GO terms in the *Tst*$^{-/-}$ liver proteome.**

**Table S3A: GO terms - Increased in ND *Tst*$^{-/-}$ liver vs ND C57Bl/6J liver**

| ID | Name | Pvalue |
|---|---|---|
| GO:0008483 | transaminase activity | 1.37E-04 |
| GO:0016769 | transferase activity, transferring nitrogenous groups | 1.37E-04 |
| GO:0005739 | mitochondrion | 1.15E-03 |
| GO:0030170 | pyridoxal phosphate binding | 2.33E-03 |
| GO:0006520 | cellular amino acid metabolic process | 2.45E-03 |
| GO:1901605 | alpha-amino acid metabolic process | 2.99E-03 |
| GO:0035970 | peptidyl-threonine dephosphorylation | 3.57E-03 |
| GO:0042347 | negative regulation of NF-kappaB import into nucleus | 3.57E-03 |
| GO:0004021 | L-alanine:2-oxoglutarate aminotransferase activity | 3.65E-03 |
| GO:0047635 | alanine-oxo-acid transaminase activity | 3.65E-03 |
| GO:0003824 | catalytic activity | 3.85E-03 |
| GO:0051213 | dioxygenase activity | 5.96E-03 |
| GO:0045893 | positive regulation of transcription, DNA-templated | 7.32E-03 |
| GO:1902680 | positive regulation of RNA biosynthetic process | 8.29E-03 |
| GO:0000959 | mitochondrial RNA metabolic process | 1.03E-02 |
| GO:0042851 | L-alanine metabolic process | 1.03E-02 |
| GO:0006522 | alanine metabolic process | 1.03E-02 |
| GO:0009078 | pyruvate family amino acid metabolic process | 1.03E-02 |
| GO:0006304 | DNA modification | 1.03E-02 |
| GO:0042345 | regulation of NF-kappaB import into nucleus | 1.03E-02 |
| GO:0042348 | NF-kappaB import into nucleus | 1.03E-02 |
| GO:0048524 | positive regulation of viral process | 1.03E-02 |
| GO:0043902 | positive regulation of multi-organism process | 1.03E-02 |
| GO:0004407 | histone deacetylase activity | 1.05E-02 |
| GO:0033558 | protein deacetylase activity | 1.05E-02 |
| GO:0051254 | positive regulation of RNA metabolic process | 1.05E-02 |
| GO:0032259 | methylation | 1.20E-02 |
| GO:0030111 | regulation of Wnt signaling pathway | 1.35E-02 |
| GO:0050714 | positive regulation of protein secretion | 1.35E-02 |
| GO:0010557 | positive regulation of macromolecule biosynthetic process | 1.47E-02 |
| GO:0000275 | mitochondrial proton-transporting ATP synthase complex, catalytic core F(1) | 1.88E-02 |
| GO:0045261 | proton-transporting ATP synthase complex, catalytic core F(1) | 1.88E-02 |
| GO:0005761 | mitochondrial ribosome | 1.88E-02 |
| GO:0000313 | organellar ribosome | 1.88E-02 |
| GO:0009063 | cellular amino acid catabolic process | 1.96E-02 |
| GO:1901607 | alpha-amino acid biosynthetic process | 1.98E-02 |
| GO:0006499 | N-terminal protein myristoylation | 1.98E-02 |
| GO:0030177 | positive regulation of Wnt signaling pathway | 1.98E-02 |
| GO:0006498 | N-terminal protein lipidation | 1.98E-02 |
| GO:0018377 | protein myristoylation | 1.98E-02 |
| GO:0042992 | negative regulation of transcription factor import into nucleus | 1.98E-02 |
| GO:0042308 | negative regulation of protein import into nucleus | 1.98E-02 |
| GO:0070403 | NAD+ binding | 2.02E-02 |

| | | |
|---|---|---|
| GO:0019213 | deacetylase activity | 2.02E-02 |
| GO:0016740 | transferase activity | 2.04E-02 |
| GO:0009891 | positive regulation of biosynthetic process | 2.17E-02 |
| GO:0006732 | coenzyme metabolic process | 2.33E-02 |
| GO:0016055 | Wnt signaling pathway | 2.36E-02 |
| GO:0042157 | lipoprotein metabolic process | 2.44E-02 |
| GO:0006544 | glycine metabolic process | 2.44E-02 |
| GO:0019752 | carboxylic acid metabolic process | 2.48E-02 |
| GO:0009108 | coenzyme biosynthetic process | 2.60E-02 |
| GO:0010628 | positive regulation of gene expression | 2.63E-02 |
| GO:0051173 | positive regulation of nitrogen compound metabolic process | 2.72E-02 |
| GO:0046394 | carboxylic acid biosynthetic process | 2.73E-02 |
| GO:0016053 | organic acid biosynthetic process | 2.73E-02 |
| GO:0043436 | oxoacid metabolic process | 2.79E-02 |
| GO:0009069 | serine family amino acid metabolic process | 2.82E-02 |
| GO:0044249 | cellular biosynthetic process | 2.85E-02 |
| GO:1901606 | alpha-amino acid catabolic process | 2.92E-02 |
| GO:0031328 | positive regulation of cellular biosynthetic process | 2.95E-02 |
| GO:0006082 | organic acid metabolic process | 3.14E-02 |
| GO:0060828 | regulation of canonical Wnt signaling pathway | 3.17E-02 |
| GO:0006497 | protein lipidation | 3.17E-02 |
| GO:0031365 | N-terminal protein amino acid modification | 3.17E-02 |
| GO:0016811 | hydrolase activity, acting on carbon-nitrogen (but not peptide) bonds, in linear amides | 3.19E-02 |
| GO:0048037 | cofactor binding | 3.23E-02 |
| GO:0004722 | protein serine/threonine phosphatase activity | 3.23E-02 |
| GO:0045935 | positive regulation of nucleobase-containing compound metabolic process | 3.41E-02 |
| GO:0045202 | synapse | 3.44E-02 |
| GO:0051186 | cofactor metabolic process | 3.55E-02 |
| GO:1901564 | organonitrogen compound metabolic process | 3.82E-02 |
| GO:0007178 | transmembrane receptor protein serine/threonine kinase signaling pathway | 3.87E-02 |
| GO:0050708 | regulation of protein secretion | 3.87E-02 |
| GO:0009058 | biosynthetic process | 3.94E-02 |
| GO:0008652 | cellular amino acid biosynthetic process | 4.37E-02 |
| GO:0043650 | dicarboxylic acid biosynthetic process | 4.57E-02 |
| GO:0035601 | protein deacylation | 4.57E-02 |
| GO:0098732 | macromolecule deacylation | 4.57E-02 |
| GO:0090092 | regulation of transmembrane receptor protein serine/threonine kinase signaling pathway | 4.57E-02 |
| GO:0007179 | transforming growth factor beta receptor signaling pathway | 4.57E-02 |
| GO:0006172 | ADP biosynthetic process | 4.57E-02 |
| GO:0009180 | purine ribonucleoside diphosphate biosynthetic process | 4.57E-02 |
| GO:0046031 | ADP metabolic process | 4.57E-02 |
| GO:0009136 | purine nucleoside diphosphate biosynthetic process | 4.57E-02 |
| GO:0009179 | purine ribonucleoside diphosphate metabolic process | 4.57E-02 |
| GO:0009135 | purine nucleoside diphosphate metabolic process | 4.57E-02 |
| GO:0046961 | proton-transporting ATPase activity, rotational mechanism | 4.66E-02 |
| GO:0046933 | proton-transporting ATP synthase activity, rotational mechanism | 4.66E-02 |
| GO:0036442 | hydrogen-exporting ATPase activity | 4.66E-02 |
| GO:0005546 | phosphatidylinositol-4,5-bisphosphate binding | 4.66E-02 |

| | | |
|---|---|---|
| GO:1902936 | phosphatidylinositol bisphosphate binding | 4.66E-02 |
| GO:0001085 | RNA polymerase II transcription factor binding | 4.66E-02 |
| GO:0047617 | acyl-CoA hydrolase activity | 4.66E-02 |
| GO:0042558 | pteridine-containing compound metabolic process | 4.72E-02 |

**Table S3B: GO terms - Decreased in ND *Tst*$^{-/-}$ liver**

| ID | Name | Pvalue |
|---|---|---|
| GO:0004745 | retinol dehydrogenase activity | 8.89E-06 |
| GO:0004364 | glutathione transferase activity | 9.72E-06 |
| GO:0043295 | glutathione binding | 4.97E-05 |
| GO:1900750 | oligopeptide binding | 4.97E-05 |
| GO:0016765 | transferase activity, transferring alkyl or aryl (other than methyl) groups | 1.10E-04 |
| GO:0032564 | dATP binding | 1.66E-04 |
| GO:0032558 | adenyl deoxyribonucleotide binding | 1.66E-04 |
| GO:0005783 | endoplasmic reticulum | 4.81E-04 |
| GO:0032554 | purine deoxyribonucleotide binding | 6.36E-04 |
| GO:0032552 | deoxyribonucleotide binding | 6.36E-04 |
| GO:0030855 | epithelial cell differentiation | 8.42E-04 |
| GO:0012505 | endomembrane system | 1.13E-03 |
| GO:0006749 | glutathione metabolic process | 1.31E-03 |
| GO:0019904 | protein domain specific binding | 1.43E-03 |
| GO:0072341 | modified amino acid binding | 2.27E-03 |
| GO:0065008 | regulation of biological quality | 2.71E-03 |
| GO:0070887 | cellular response to chemical stimulus | 2.83E-03 |
| GO:0005102 | receptor binding | 2.88E-03 |
| GO:0004190 | aspartic-type endopeptidase activity | 3.05E-03 |
| GO:0070001 | aspartic-type peptidase activity | 3.05E-03 |
| GO:0002135 | CTP binding | 3.05E-03 |
| GO:0017098 | sulfonylurea receptor binding | 3.05E-03 |
| GO:0007269 | neurotransmitter secretion | 3.13E-03 |
| GO:0006635 | fatty acid beta-oxidation | 3.19E-03 |
| GO:0051640 | organelle localization | 3.19E-03 |
| GO:0008306 | associative learning | 3.19E-03 |
| GO:0018916 | nitrobenzene metabolic process | 3.19E-03 |
| GO:0070458 | cellular detoxification of nitrogen compound | 3.19E-03 |
| GO:0051410 | detoxification of nitrogen compound | 3.19E-03 |
| GO:2000301 | negative regulation of synaptic vesicle exocytosis | 3.19E-03 |
| GO:0032486 | Rap protein signal transduction | 3.19E-03 |
| GO:1902804 | negative regulation of synaptic vesicle transport | 3.19E-03 |
| GO:0045793 | positive regulation of cell size | 3.19E-03 |
| GO:0090316 | positive regulation of intracellular protein transport | 4.69E-03 |
| GO:0042221 | response to chemical | 4.99E-03 |
| GO:1901214 | regulation of neuron death | 5.57E-03 |
| GO:0051234 | establishment of localization | 6.32E-03 |
| GO:0044708 | single-organism behavior | 6.45E-03 |

| | | |
|---|---|---|
| GO:0070997 | neuron death | 6.60E-03 |
| GO:0071310 | cellular response to organic substance | 6.92E-03 |
| GO:0016757 | transferase activity, transferring glycosyl groups | 6.93E-03 |
| GO:1901681 | sulfur compound binding | 7.00E-03 |
| GO:0016758 | transferase activity, transferring hexosyl groups | 7.14E-03 |
| GO:0010033 | response to organic substance | 7.61E-03 |
| GO:0031526 | brush border membrane | 7.71E-03 |
| GO:0006810 | transport | 8.35E-03 |
| GO:0006629 | lipid metabolic process | 8.67E-03 |
| GO:0005550 | pheromone binding | 8.82E-03 |
| GO:0005549 | odorant binding | 8.82E-03 |
| GO:0004579 | dolichyl-diphosphooligosaccharide-protein glycotransferase activity | 8.82E-03 |
| GO:0002134 | UTP binding | 8.82E-03 |
| GO:0032551 | pyrimidine ribonucleoside binding | 8.82E-03 |
| GO:0001884 | pyrimidine nucleoside binding | 8.82E-03 |
| GO:0006518 | peptide metabolic process | 9.05E-03 |
| GO:0035690 | cellular response to drug | 9.23E-03 |
| GO:0051883 | killing of cells in other organism involved in symbiotic interaction | 9.23E-03 |
| GO:0051818 | disruption of cells of other organism involved in symbiotic interaction | 9.23E-03 |
| GO:0044364 | disruption of cells of other organism | 9.23E-03 |
| GO:0031640 | killing of cells of other organism | 9.23E-03 |
| GO:0045920 | negative regulation of exocytosis | 9.23E-03 |
| GO:0033160 | positive regulation of protein import into nucleus, translocation | 9.23E-03 |
| GO:0006982 | response to lipid hydroperoxide | 9.23E-03 |
| GO:0051769 | regulation of nitric-oxide synthase biosynthetic process | 9.23E-03 |
| GO:0051767 | nitric-oxide synthase biosynthetic process | 9.23E-03 |
| GO:0051656 | establishment of organelle localization | 9.51E-03 |
| GO:0044765 | single-organism transport | 9.95E-03 |
| GO:0009636 | response to toxic substance | 1.05E-02 |
| GO:0002064 | epithelial cell development | 1.05E-02 |
| GO:0019395 | fatty acid oxidation | 1.05E-02 |
| GO:0034440 | lipid oxidation | 1.05E-02 |
| GO:0044439 | peroxisomal part | 1.06E-02 |
| GO:0044438 | microbody part | 1.06E-02 |
| GO:0043523 | regulation of neuron apoptotic process | 1.14E-02 |
| GO:0000038 | very long-chain fatty acid metabolic process | 1.16E-02 |
| GO:0048489 | synaptic vesicle transport | 1.16E-02 |
| GO:0097480 | establishment of synaptic vesicle localization | 1.16E-02 |
| GO:0097479 | synaptic vesicle localization | 1.16E-02 |
| GO:0036473 | cell death in response to oxidative stress | 1.16E-02 |
| GO:0001885 | endothelial cell development | 1.16E-02 |
| GO:0044242 | cellular lipid catabolic process | 1.16E-02 |
| GO:0016042 | lipid catabolic process | 1.18E-02 |
| GO:0009062 | fatty acid catabolic process | 1.21E-02 |
| GO:0005788 | endoplasmic reticulum lumen | 1.24E-02 |
| GO:0005778 | peroxisomal membrane | 1.24E-02 |
| GO:0031903 | microbody membrane | 1.24E-02 |
| GO:0042277 | peptide binding | 1.25E-02 |

| | | |
|---|---|---|
| GO:0051402 | neuron apoptotic process | 1.35E-02 |
| GO:0042803 | protein homodimerization activity | 1.35E-02 |
| GO:0016860 | intramolecular oxidoreductase activity | 1.49E-02 |
| GO:0019207 | kinase regulator activity | 1.49E-02 |
| GO:0048306 | calcium-dependent protein binding | 1.49E-02 |
| GO:0001906 | cell killing | 1.59E-02 |
| GO:0009813 | flavonoid biosynthetic process | 1.59E-02 |
| GO:0052696 | flavonoid glucuronidation | 1.59E-02 |
| GO:0052695 | cellular glucuronidation | 1.59E-02 |
| GO:0019585 | glucuronate metabolic process | 1.59E-02 |
| GO:0006063 | uronic acid metabolic process | 1.59E-02 |
| GO:0006836 | neurotransmitter transport | 1.59E-02 |
| GO:0045446 | endothelial cell differentiation | 1.59E-02 |
| GO:0003158 | endothelium development | 1.59E-02 |
| GO:0034185 | apolipoprotein binding | 1.70E-02 |
| GO:0030971 | receptor tyrosine kinase binding | 1.70E-02 |
| GO:0032557 | pyrimidine ribonucleotide binding | 1.70E-02 |
| GO:0019103 | pyrimidine nucleotide binding | 1.70E-02 |
| GO:0005777 | peroxisome | 1.77E-02 |
| GO:0042579 | microbody | 1.77E-02 |
| GO:0042760 | very long-chain fatty acid catabolic process | 1.78E-02 |
| GO:0042573 | retinoic acid metabolic process | 1.78E-02 |
| GO:0018279 | protein N-linked glycosylation via asparagine | 1.78E-02 |
| GO:0018196 | peptidyl-asparagine modification | 1.78E-02 |
| GO:0007097 | nuclear migration | 1.78E-02 |
| GO:0040023 | establishment of nucleus localization | 1.78E-02 |
| GO:2000300 | regulation of synaptic vesicle exocytosis | 1.78E-02 |
| GO:1902803 | regulation of synaptic vesicle transport | 1.78E-02 |
| GO:0046928 | regulation of neurotransmitter secretion | 1.78E-02 |
| GO:0051588 | regulation of neurotransmitter transport | 1.78E-02 |
| GO:0033158 | regulation of protein import into nucleus, translocation | 1.78E-02 |
| GO:0033194 | response to hydroperoxide | 1.78E-02 |
| GO:0051702 | interaction with symbiont | 1.78E-02 |
| GO:0070664 | negative regulation of leukocyte proliferation | 1.78E-02 |
| GO:0007268 | synaptic transmission | 1.80E-02 |
| GO:0060627 | regulation of vesicle-mediated transport | 1.80E-02 |
| GO:0031346 | positive regulation of cell projection organization | 1.96E-02 |
| GO:0071407 | cellular response to organic cyclic compound | 1.96E-02 |
| GO:0032388 | positive regulation of intracellular transport | 1.96E-02 |
| GO:0033554 | cellular response to stress | 1.97E-02 |
| GO:0043178 | alcohol binding | 1.97E-02 |
| GO:0045859 | regulation of protein kinase activity | 2.03E-02 |
| GO:0033218 | amide binding | 2.05E-02 |
| GO:0017157 | regulation of exocytosis | 2.10E-02 |
| GO:0009812 | flavonoid metabolic process | 2.10E-02 |
| GO:0071495 | cellular response to endogenous stimulus | 2.25E-02 |
| GO:0051179 | localization | 2.26E-02 |
| GO:0007264 | small GTPase mediated signal transduction | 2.28E-02 |

| | | |
|---|---|---|
| GO:0009617 | response to bacterium | 2.34E-02 |
| GO:0050896 | response to stimulus | 2.46E-02 |
| GO:0015020 | glucuronosyltransferase activity | 2.53E-02 |
| GO:0072329 | monocarboxylic acid catabolic process | 2.55E-02 |
| GO:0005903 | brush border | 2.57E-02 |
| GO:0007611 | learning or memory | 2.68E-02 |
| GO:0004576 | oligosaccharyl transferase activity | 2.73E-02 |
| GO:0016937 | short-branched-chain-acyl-CoA dehydrogenase activity | 2.73E-02 |
| GO:0016290 | palmitoyl-CoA hydrolase activity | 2.73E-02 |
| GO:0045454 | cell redox homeostasis | 2.77E-02 |
| GO:0000302 | response to reactive oxygen species | 2.77E-02 |
| GO:0007610 | behavior | 2.78E-02 |
| GO:0030258 | lipid modification | 2.84E-02 |
| GO:0032386 | regulation of intracellular transport | 2.84E-02 |
| GO:0010592 | positive regulation of lamellipodium assembly | 2.85E-02 |
| GO:0010591 | regulation of lamellipodium assembly | 2.85E-02 |
| GO:1902745 | positive regulation of lamellipodium organization | 2.85E-02 |
| GO:1902743 | regulation of lamellipodium organization | 2.85E-02 |
| GO:0051409 | response to nitrosative stress | 2.85E-02 |
| GO:0072384 | organelle transport along microtubule | 2.85E-02 |
| GO:0048747 | muscle fiber development | 2.85E-02 |
| GO:0051817 | modification of morphology or physiology of other organism involved in symbiotic interaction | 2.85E-02 |
| GO:0035821 | modification of morphology or physiology of other organism | 2.85E-02 |
| GO:0071320 | cellular response to cAMP | 2.85E-02 |
| GO:0016079 | synaptic vesicle exocytosis | 2.85E-02 |
| GO:0051049 | regulation of transport | 2.95E-02 |
| GO:0051649 | establishment of localization in cell | 2.98E-02 |
| GO:0019899 | enzyme binding | 2.99E-02 |
| GO:0060429 | epithelium development | 2.99E-02 |
| GO:0003995 | acyl-CoA dehydrogenase activity | 3.15E-02 |
| GO:0005215 | transporter activity | 3.24E-02 |
| GO:0051641 | cellular localization | 3.31E-02 |
| GO:0070372 | regulation of ERK1 and ERK2 cascade | 3.35E-02 |
| GO:0051716 | cellular response to stimulus | 3.38E-02 |
| GO:0031406 | carboxylic acid binding | 3.62E-02 |
| GO:0051222 | positive regulation of protein transport | 3.76E-02 |
| GO:0003824 | catalytic activity | 3.77E-02 |
| GO:0016597 | amino acid binding | 3.84E-02 |
| GO:0043549 | regulation of kinase activity | 3.85E-02 |
| GO:1901701 | cellular response to oxygen-containing compound | 3.89E-02 |
| GO:0043177 | organic acid binding | 3.90E-02 |
| GO:0004860 | protein kinase inhibitor activity | 3.95E-02 |
| GO:0019210 | kinase inhibitor activity | 3.95E-02 |
| GO:0004602 | glutathione peroxidase activity | 3.95E-02 |
| GO:0047617 | acyl-CoA hydrolase activity | 3.95E-02 |
| GO:0008144 | drug binding | 4.01E-02 |
| GO:0046983 | protein dimerization activity | 4.04E-02 |
| GO:0044432 | endoplasmic reticulum part | 4.04E-02 |

| | | |
|---|---|---|
| GO:0050890 | cognition | 4.09E-02 |
| GO:0051650 | establishment of vesicle localization | 4.09E-02 |
| GO:0051648 | vesicle localization | 4.09E-02 |
| GO:0014074 | response to purine-containing compound | 4.09E-02 |
| GO:0019901 | protein kinase binding | 4.11E-02 |
| GO:0042026 | protein refolding | 4.12E-02 |
| GO:0007612 | learning | 4.12E-02 |
| GO:1900408 | negative regulation of cellular response to oxidative stress | 4.12E-02 |
| GO:1902883 | negative regulation of response to oxidative stress | 4.12E-02 |
| GO:0043525 | positive regulation of neuron apoptotic process | 4.12E-02 |
| GO:0010976 | positive regulation of neuron projection development | 4.12E-02 |
| GO:0061028 | establishment of endothelial barrier | 4.12E-02 |
| GO:0000060 | protein import into nucleus, translocation | 4.12E-02 |
| GO:0001836 | release of cytochrome c from mitochondria | 4.12E-02 |
| GO:0043603 | cellular amide metabolic process | 4.23E-02 |
| GO:0051338 | regulation of transferase activity | 4.23E-02 |
| GO:0070201 | regulation of establishment of protein localization | 4.23E-02 |
| GO:0071417 | cellular response to organonitrogen compound | 4.32E-02 |
| GO:0033157 | regulation of intracellular protein transport | 4.32E-02 |
| GO:0051050 | positive regulation of transport | 4.39E-02 |
| GO:0016192 | vesicle-mediated transport | 4.53E-02 |
| GO:0051082 | unfolded protein binding | 4.58E-02 |
| GO:0008194 | UDP-glycosyltransferase activity | 4.63E-02 |
| GO:0005496 | steroid binding | 4.63E-02 |
| GO:0006457 | protein folding | 4.64E-02 |
| GO:0060341 | regulation of cellular localization | 4.71E-02 |
| GO:0045184 | establishment of protein localization | 4.90E-02 |
| GO:0045926 | negative regulation of growth | 4.91E-02 |
| GO:0001505 | regulation of neurotransmitter levels | 4.91E-02 |
| GO:1901699 | cellular response to nitrogen compound | 4.93E-02 |
| GO:0097193 | intrinsic apoptotic signaling pathway | 4.93E-02 |

**Table S3. Tst Deletion results in differential hepatic protein abundance of GO terms.** (**A**) Significant GO terms represented by proteins that are more abundant in the ND-fed *Tst*$^{-/-}$ liver compared with normal diet-fed C57Bl/6J. (**B**) Significant GO terms represented by proteins that are less abundant in the ND-fed *Tst*$^{-/-}$ liver compared with normal diet-fed C57Bl/6J. '**Genes**' indicates the number of genes in the *Tst*$^{-/-}$ that represent the changes driving the GO term. The lists include all regulated GO terms at a significance threshold of P < 0.05

| Table S4. Sulfide metabolism proteins in liver proteome (*Tst*$^{-/-}$ vs C57Bl/6J, ND-fed) | | | |
|---|---|---|---|
| **Feature ID** | **Name** | **Fold Change** | **Significance** |
| Q3UW66 | MPST Mercaptopyruvate sulfurtrasferase | 1.27 | ** |
| Q8R086 | SUOX – Sulfite Oxidase | 1.06 | Ns |
| Q3UDS4 | SQOR – Sulfide quinone reductase-like | 1.06 | Ns |
| Q91WT9 | CBS – Cystathionine beta-synthase | 1.03 | Ns |
| Q9DCM0 | ETHE1 – Ethylmalonic encephalopathy 1 | -1.01 | Ns |
| Q8VCNS | CTH – Cystathionine gamma-lyase | -1.02 | Ns |
| * Raw P < 0.05, ** Adjusted P < 0.05 | | | |

**Table S4. *Tst* Deletion selectively regulates MPST in the sulfide pathway of normal diet–fed mice.** Relative peptide abundance of proteins of the sulfide production and disposal pathway from the liver proteome of normal diet (ND) fed mice . '**Fold Change**' indicates the relative abundance of the protein in *Tst*$^{-/-}$ relative to C57Bl/6J.

| Table S5. GO terms - Nutrient metabolism (*Tst*$^{-/-}$ vs C57Bl/6J liver, ND-fed) | | | |
|---|---|---|---|
| **GO-ID** | **Name** | **Genes** | **Significance** |
| **Reduced in ND *Tst*$^{-/-}$ liver** | | | |
| 0006629 | Lipid metabolic process | 19 | ** |
| 0006631 | Fatty acid beta-oxidation | 7 | ** |
| 0003995 | Acyl-CoA dehydrogenase activity | 3 | * |
| 0047617 | Acyl-CoA hydrolase activity | 2 | * |
| * P < 0.05, ** P < 0.01 | | | |

**Table S5. Tst Deletion results in reduction of selective fatty acid specific GO terms.** Significant GO terms (glucose or lipid related) represented by proteins that are less abundant in the ND-fed *Tst*$^{-/-}$ liver compared with ND-fed C57Bl/6J. '**Genes**' indicates the number of genes in the *Tst*$^{-/-}$ that represent the changes driving the GO term.

| Table S6 Insulin regulated proteins in normal diet fed *Tst*$^{-/-}$ and C57Bl/6J | | | |
|---|---|---|---|
| **(A) Abundance of peptides of insulin–induced proteins (*Tst*$^{-/-}$ vs C57Bl/6J, ND-fed)** | | | |
| **Feature ID** | **Name** | **Fold change** | **Significance** |
| Q3UGT1 | CPT1A | 1.03 | Ns |
| P19096 | FASN | -1.04 | Ns |
| Q3UDA8 | CPT2 | -1.05 | Ns |
| Q3V2G1 | APOA1 | -1.10 | Ns |
| Q5SVI5 | GCK | -1.14 | * |
| **(B) Abundance of peptides of insulin–suppressed proteins (*Tst*$^{-/-}$ vs C57Bl/6J, ND-fed)** | | | |
| **Feature ID** | **Name** | **Fold change** | **Significance** |
| QO5421 | CYP2E1 | 1.13 | ** |
| Q9D6M3 | SLC25AA2 | -1.03 | Ns |
| Q8CI37 | PCK1 | -1.09 | Ns |
| Q3UJ70 | HMGCS1 | -1.09 | Ns |
| O08601 | MTTP | -1.20 | ** |
| *** Raw P < 0.05, ** Adjusted P < 0.05** | | | |

**Table S6. Proteins regulated by insulin are broadly comparable in expression between *Tst*$^{-/-}$ and C57Bl/6J on normal diet.** Relative abundance in proteins that are known to be induced (A) or suppressed (B) by insulin in the liver, from the liver proteome of normal diet fed mice. '**Fold Change**' indicates the relative abundance of the protein in *Tst*$^{-/-}$ relative to C57Bl/6J.

| Table S7 Insulin regulated proteins in high fat diet fed *Tst*$^{-/-}$ and C57Bl/6J | | | |
|---|---|---|---|
| **(A) Abundance of peptides of insulin–induced proteins (*Tst*$^{-/-}$ vs C57Bl/6J, ND-fed)** | | | |
| **Feature ID** | **Name** | **Fold change** | **Significance** |
| Q3UGT1 | CPT1A | 1.03 | Ns |
| P19096 | FASN | -1.04 | Ns |
| Q3UDA8 | CPT2 | -1.05 | Ns |
| Q3V2G1 | APOA1 | -1.1 | Ns |
| Q5SVI5 | GCK | -1.14 | * |
| **(B) Abundance of peptides of insulin–suppressed proteins (*Tst*$^{-/-}$ vs C57Bl/6J, ND-fed)** | | | |
| **Feature ID** | **Name** | **Fold change** | **Significance** |
| QO5421 | CYP2E1 | -1.05 | Ns |
| Q9D6M3 | SLC25AA2 | -1.01 | Ns |
| Q8CI37 | PCK1 | 1.09 | Ns |
| Q3UJ70 | HMGCS1 | -1.04 | Ns |
| O08601 | MTTP | 1.09 | * |
| *** Raw P < 0.05, ** Adjusted P < 0.05** | | | |

**Table S7. Proteins regulated by insulin are broadly comparable in expression between *Tst*$^{-/-}$ and C57Bl/6J on high fat diet.** Relative abundance in proteins that are known to be induced (A) or suppressed (B) by insulin in the liver, from the liver proteome of high fat diet fed mice. '**Fold Change**' indicates the relative abundance of the protein in *Tst*$^{-/-}$ relative to C57Bl/6J.

| Table S8. KEGG pathways shared by high fat feeding and TST deletion | | | |
|---|---|---|---|
| **Entry** | **Name** | **Comparison** | **Significance** |
| **A. Shared up-regulated pathways** | | | |
| 00260 | Glycine, serine and threonine metabolism | *Tst*$^{-/-}$ vs 6J | ** |
| | | HFD vs ND | ** |
| **B. Shared down-regulated pathways** | | | |
| 00980 | Metabolism of xenobiotics by cytochrome P450 | *Tst*$^{-/-}$ vs 6J | **** |
| | | HFD vs ND | * |
| 00982 | Drug metabolism – cytochrome P450 | *Tst*$^{-/-}$ vs 6J | **** |
| | | HFD vs ND | * |
| 04142 | Lysosome | *Tst*$^{-/-}$ vs 6J | ** |
| | | HFD vs ND | **** |
| 04390 | Hippo signaling pathway | *Tst*$^{-/-}$ vs 6J | ** |
| | | HFD vs ND | **** |
| 05215 | Prostate cancer | *Tst*$^{-/-}$ vs 6J | ** |
| | | HFD vs ND | * |
| 04024 | cAMP signaling pathway | *Tst*$^{-/-}$ vs 6J | * |
| | | HFD vs 6J | * |
| 04141 | Protein processing endoplasmic reticulum | *Tst*$^{-/-}$ vs 6J | * |
| | | HFD vs 6J | ** |
| 05211 | Renal cell carcinoma | *Tst*$^{-/-}$ vs 6J | * |
| | | HFD vs 6J | * |
| 04722 | Neurotrophin signaling pathway | *Tst*$^{-/-}$ vs 6J | * |
| | | HFD vs 6J | * |
| 04110 | Cell cycle | *Tst*$^{-/-}$ vs 6J | * |
| | | HFD vs 6J | ** |
| 04918 | Thyroid hormone synthesis | *Tst*$^{-/-}$ vs 6J | * |
| | | HFD vs 6J | *** |
| 04612 | Antigen processing and presentation | *Tst*$^{-/-}$ vs 6J | * |
| | | HFD vs 6J | ** |
| *** P < 0.05, ** P < 0.01, *** P < 0.001 **** P < 0.0001** | | | |

**Table S8. KEGG Pathways shared by high fat feeding and TST deletion.** (**A**) KEGG pathways that are significantly up-regulated in the same direction by both high fat diet (HFD vs ND), and *Tst* deletion (*Tst*$^{-/-}$ vs C57Bl/6J). (**B**) KEGG pathways that are significantly down-regulated in the same direction by both high fat diet (HFD vs ND), and *Tst* deletion (*Tst*$^{-/-}$ vs C57Bl/6J). 'Comparison' indicates the two groups being compared.

| Table S9. Effect of high fat feeding on sulfide pathway proteins (High fat diet vs ND-fed) | | | | | |
|---|---|---|---|---|---|
| **Feature ID** | **Name** | **Fold change in 6J** | **Significance** | **Fold change in *Tst*$^{-/-}$** | **Significance** |
| Q3UW66 | MPST | 1.35 | ** | 1.15 | * |
| Q8R086 | SUOX | 1.21 | ** | 1.23 | ** |
| Q545S0 | TST | 1.19 | Ns | n/a | n/a |
| Q8VCNS | CTH | -1.04 | Ns | -1.06 | Ns |
| Q9DCM0 | ETHE1 | -1.05 | Ns | 1.03 | Ns |
| Q3UDS4 | SQOR | -1.08 | Ns | 1.01 | Ns |
| Q91WT9 | CBS | -1.10 | * | -1.06 | Ns |
| *** Raw P < 0.05, ** Adjusted P < 0.05** | | | | | |

**Table S9. Effect of high fat feeding on the sulfide pathway of C57Bl/6J and *Tst*$^{-/-}$ mice.** Protein abundances of the sulfide production and disposal pathway from the liver proteome of C57Bl/6J and *Tst*$^{-/-}$ mice. '**Fold Change**' indicates the relative abundance of the protein in high fat diet fed mice relative to normal diet fed mice, shown separately for each genotype.

| Table S10. Pathways in *Tst*$^{-/-}$ that are regulated oppositely to high fat feeding | | | |
|---|---|---|---|
| **Entry** | **Name** | **Comparison** | **Direction** |
| **A** | **KEGG Pathways** | | |
| 00980 | Metabolism of xenobiotics by cytochrome P450 | *Tst*$^{-/-}$ vs 6J | Decreased |
| | | HFD vs ND | Increased |
| 00983 | Drug metabolism – other enzymes | *Tst*$^{-/-}$ vs 6J | Decreased |
| | | HFD vs ND | Increased |
| 00053 | Ascorbate and aldarate metabolism | *Tst*$^{-/-}$ vs 6J | Decreased |
| | | HFD vs ND | Increased |
| 00040 | Pentose and glucoronate interconversions | *Tst*$^{-/-}$ vs 6J | Decreased |
| | | HFD vs ND | Increased |
| 00830 | Retinol metabolism | *Tst*$^{-/-}$ vs 6J | Decreased |
| | | HFD vs ND | Increased |
| **B** | **GO Terms** | | |
| 0006629 | Lipid metabolic process | *Tst*$^{-/-}$ vs 6J | Decreased |
| | | HFD vs ND | Increased |
| 0006631 | Fatty acid beta-oxidation | *Tst*$^{-/-}$ vs 6J | Decreased |
| | | HFD vs ND | Increased |
| 0003995 | Acyl-CoA dehydrogenase activity | *Tst*$^{-/-}$ vs 6J | Decreased |
| | | HFD vs ND | Increased |
| 0047617 | Acyl-CoA hydrolase activity | *Tst*$^{-/-}$ vs 6J | Decreased |
| | | HFD vs ND | Increased |

**Table S10. KEGG pathways and GO terms that are regulated in the opposite direction by high fat feeding compared to Tst deletion.** (**A**) KEGG pathways that are regulated in the opposite direction by high fat diet (HFD-fed C57Bl/6J vs ND-fed C57Bl/6J), to *Tst* deletion (ND-fed *Tst*$^{-/-}$ vs C57Bl/6J). (**B**) GO terms that are regulated in the opposite direction by high fat diet (HFD-fed C57Bl/6J vs ND-fed C57Bl/6J), to *Tst* deletion (ND-fed *Tst*$^{-/-}$ vs C57Bl/6J). '**Comparison'** Indicates the two groups being compared. '**Direction**' indicates whether the protein abundance is decreased or increased in the first group relative to the second.

### Supplementary Figures

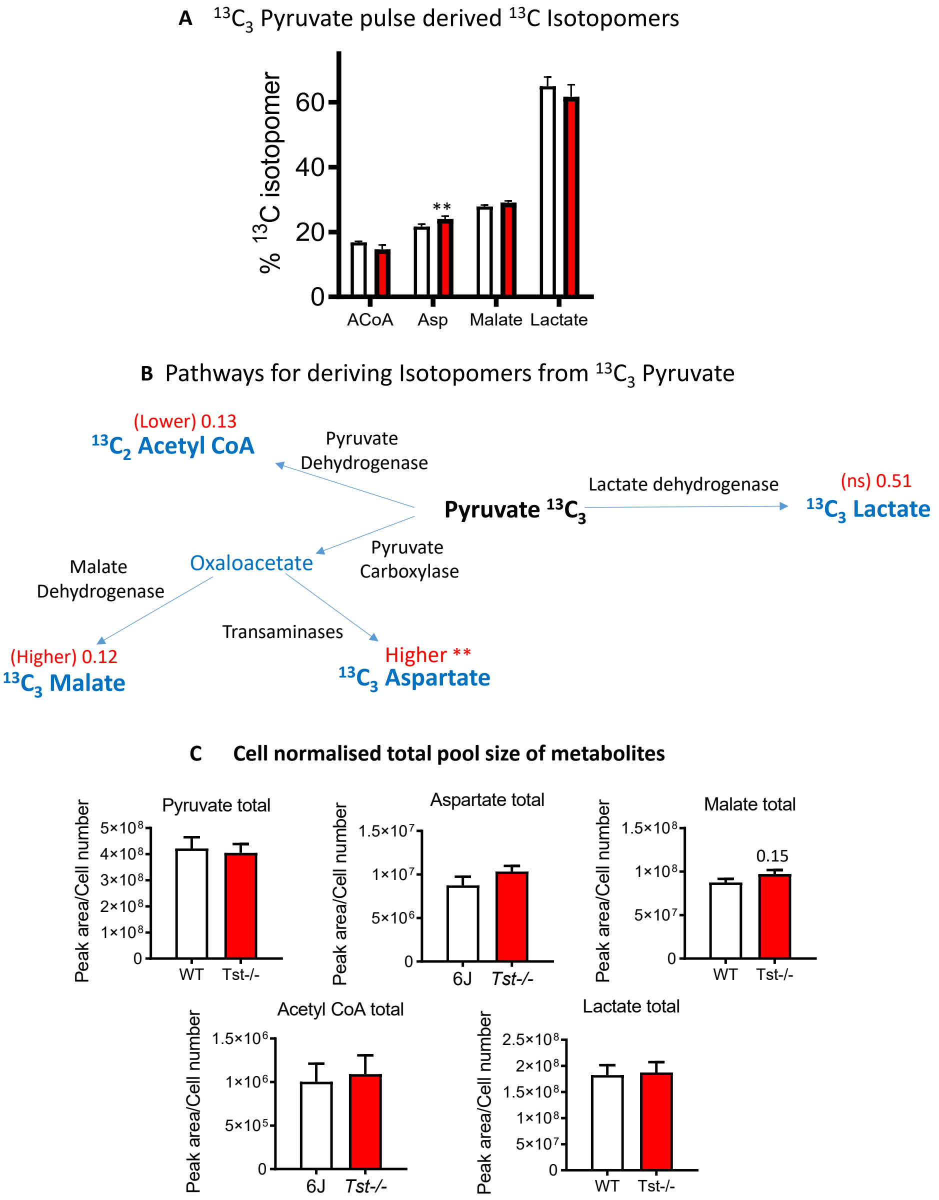


**Figure S1. Metabolic fates of $^{13}C_3$ pyruvate are altered in hepatocytes from *Tst*$^{-/-}$ mice.** **(A)** Histogram showing isotopomers derived from $^{13}C_3$ pyruvate from C57Bl/6J (white bars n=5) and *Tst*$^{-/-}$ (red bars n = 4) cultured hepatocytes. Data represents the amount of the most abundant isotopomer detected by mass spectrometry, as a percentage of the total detected metabolite (total includes unlabelled $^{12}C$ and all detectedf $^{13}C$ isotopomers). Counts were first normalised to cell number. Isotopomers shown are ACoA ($^{13}C_2$ acetyl CoA), Asp ($^{13}C_3$ aspartate), $^{13}C_3$ malate and $^{13}C_3$ lactate. **(B)** Diagram representing metabolites derived from pyruvate. Oxaloacetate was not detected, but is indicated as an intermediate to production of mallet or asparate. The isotopomer shown on the diagram represents to most abundant detected following pulse with $^{13}C_3$ pyruvate. In red is the direction of change in the *Tst*$^{-/-}$, with significance or P-values (when less than 0.2) from T-tests**. (C)** Histograms showing the total pool size of each metabolite. Data is cell normalised mass spec counts from all isotopomers of the given metabolite, including the relevant unlabelled $^{12}C$ species. Data are represented as mean ± SEM. Each metabolite was analysed using a T-test, ** indicates that $P < 0.01$. P-values less than 0.2 are also indicated for showing trends.

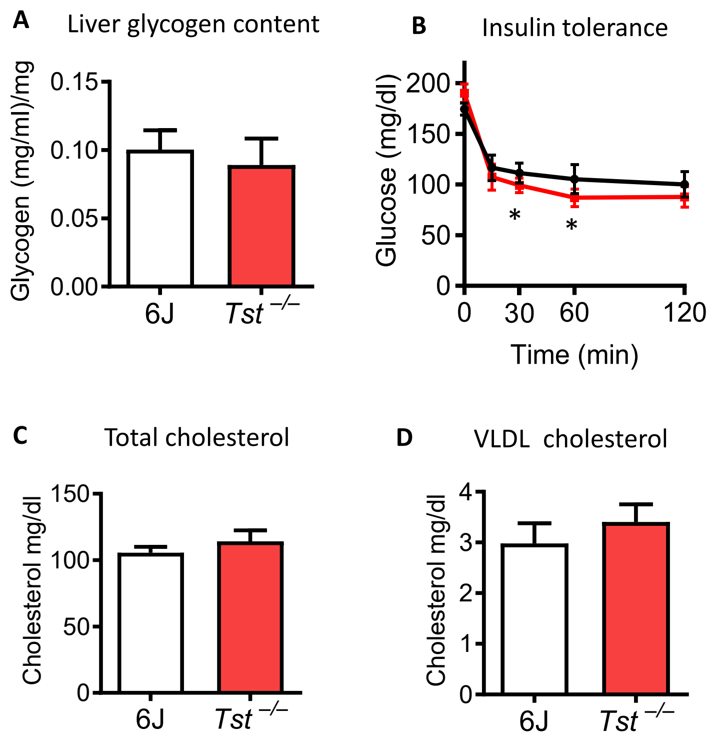


**Figure S2. Insulin–regulated metabolic parameters in liver and plasma of *Tst* $^{-/-}$ mice.** (**A**) Glycogen measured from whole liver from normal diet-fed 4 hour fasted C57Bl/6J (6J; white bar, n = 5), and *Tst*$^{-/-}$ (red bar, n = 5) mice. Data are represented as mean ±SEM. (**B**) Plasma glucose (mg/dl), over 120 minutes following insulin administration (i.p., 1mU/g) in normal diet-fed 4 hour fasted C57Bl/6J (black line, n = 8) and *Tst*$^{-/-}$ (red line, n = 7) mice. (**C**) HPLC quantified total plasma cholesterol in normal diet-fed 4 hour fasted C57Bl/6J (white bar, n = 6) and *Tst*$^{-/-}$ (red bar, n = 6) mice. (**D**) HPLC quantified VLDL plasma cholesterol in normal diet-fed 4 hour fasted C57Bl/6J (white bar, n = 6), and *Tst*$^{-/-}$ (red bar, n = 6) mice. For (**B**) a Repeated Measures analysis demonstrated a significant effect of time (****) and an interaction between time and genotype (*). T-tests revealed that the decrement of glucose from baseline at 30 and 60 minutes after insulin was greater in the *Tst*$^{-/-}$ (*).

A

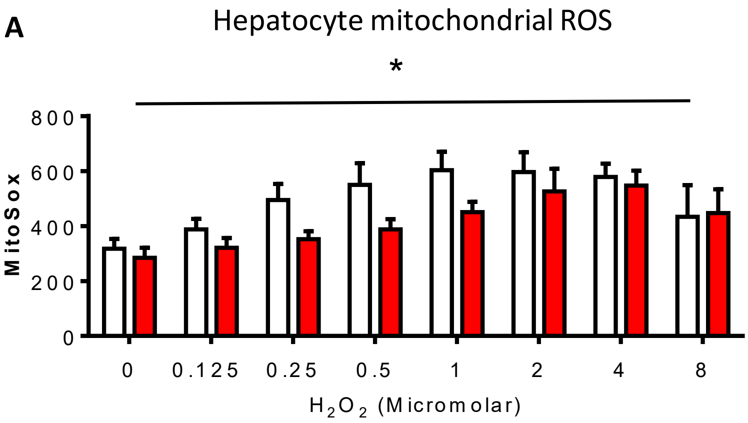


**Figure S3. Hepatocytes from *Tst* $^{-/-}$ mice resist hydrogen peroxide induced mitochondrial reactive species accumulation. (A)** Mitochondrial reactive oxygen species measured from primary hepatocytes by MitoSox fluorescence from C57Bl/6J (white bars, n = 7) and *Tst*$^{-/-}$ (red bars, n = 7). Cells were exposed to a range of doses of $H_2O_2$ prior to MitoSox incubation and fluorescent detection. Data are represented as mean ±SEM. Significance was calculated using 2-WAY ANOVA for $H_2O_2$ dose and genotype. A significant effect of genotype is represented above the histogram with a *. $H_2O_2$ was significant to $P < 0.001$ (not represented on the histogram).

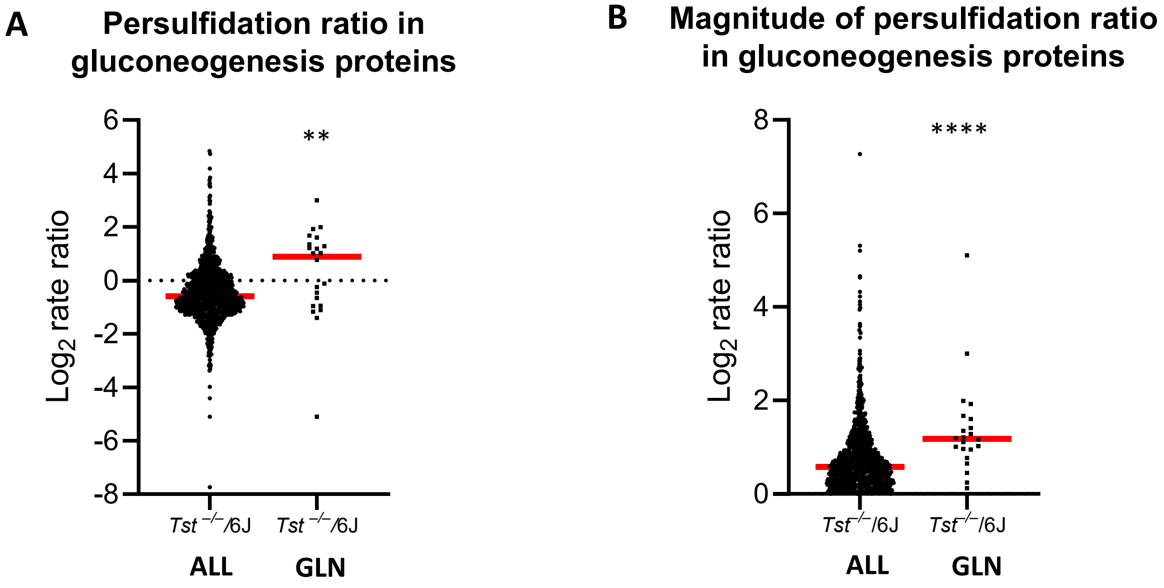


**Figure S4. Persulfidation in the gluconeogenesis pathway is significantly different to global persulfidation patterns in the liver of the *Tst*$^{-/-}$ mice.** (**A**) Beeswarm plots showing the persulfidation $\log_2$ rate ratio (*Tst*$^{-/-}$ divided by 6J) for peptides in the entire data set (ALL), alongside the $\log_2$ rate ratio for peptides corresponding to proteins of gluconeogenesis (GLN). (**B**) Beeswarm plots showing the magnitude of the $\log_2$ rate ratio (independent to direction of change), for peptides in the entire data set (ALL), alongside the $\log_2$ rate ratios for peptides corresponding to proteins of gluconeogenesis (GLN). Data are represented as individual peptide $\log_2$ rate ratio values, with the median represented as a red line. Significance was calculated using the Mann-Whitney U non parametric T-test. ** $P < 0.01$, **** $P < 0.0001$.

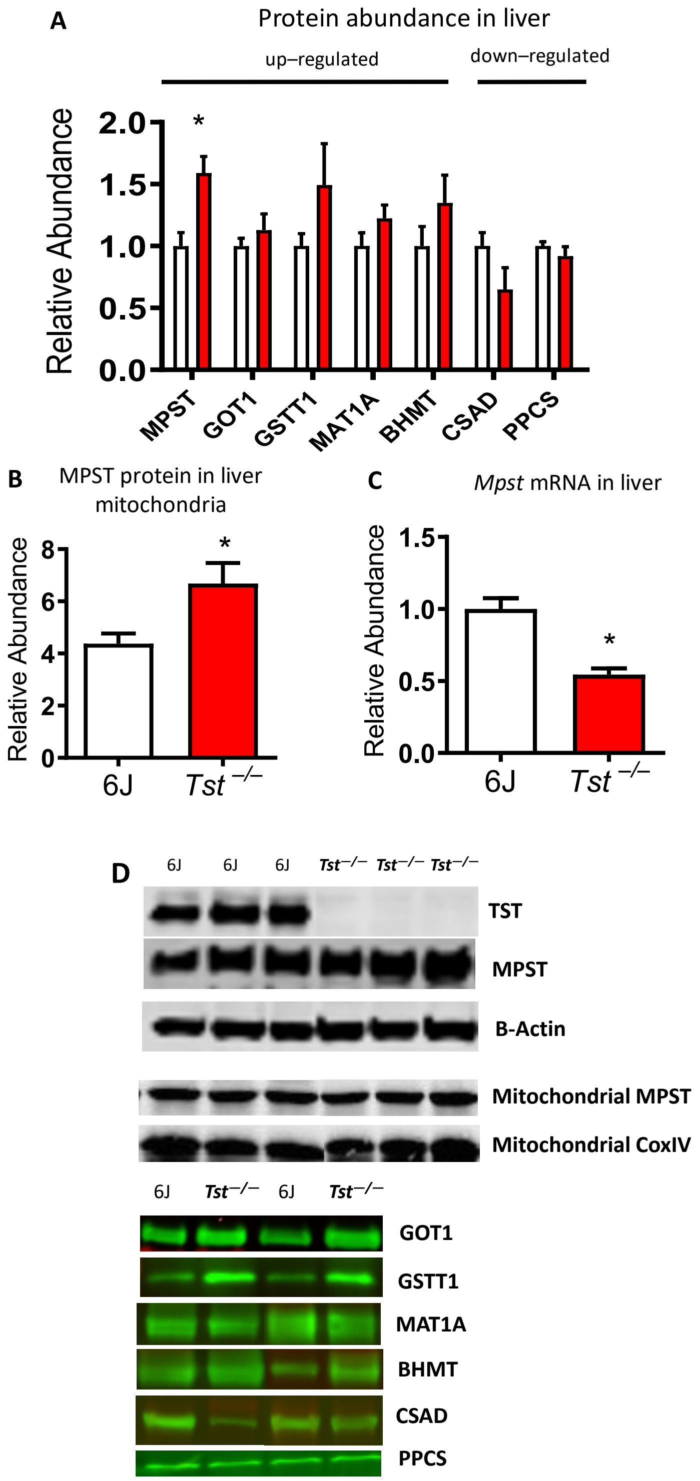


**Figure S5. Validation of proteomic profiles by select western blot is exemplified by increased mitochondrial MPST.** (**A**) Quantification of western blots for a range of proteins found significantly up or down-regulated in the liver proteome of normal diet-fed 4 hour fasted C57Bl/6J (6J; white bar, n = 4-6) and *Tst*$^{-/-}$ (red bar, n = 4-6) mice. (**B**) Quantification of western blots for MPST from isolated liver mitochondria of normal diet-fed 4 hour fasted C57Bl/6J (white bar, n = 6), and *Tst*$^{-/-}$ (red bar, n = 6) mice. (**C**) *Mpst* mRNA quantified by real time PCR from liver of normal diet-fed C57Bl/6J (6J; white bar, n = 6) and *Tst*$^{-/-}$ (red bar, n = 6) mice. (**D**) Representative blots from LICOR imaging for the data quantified in (**A**). Data are represented as mean ±SEM. Significance was calculated using un-paired two-tailed student's t-test. * $P < 0.05$, ** $P < 0.01$, *** $P < 0.001$.

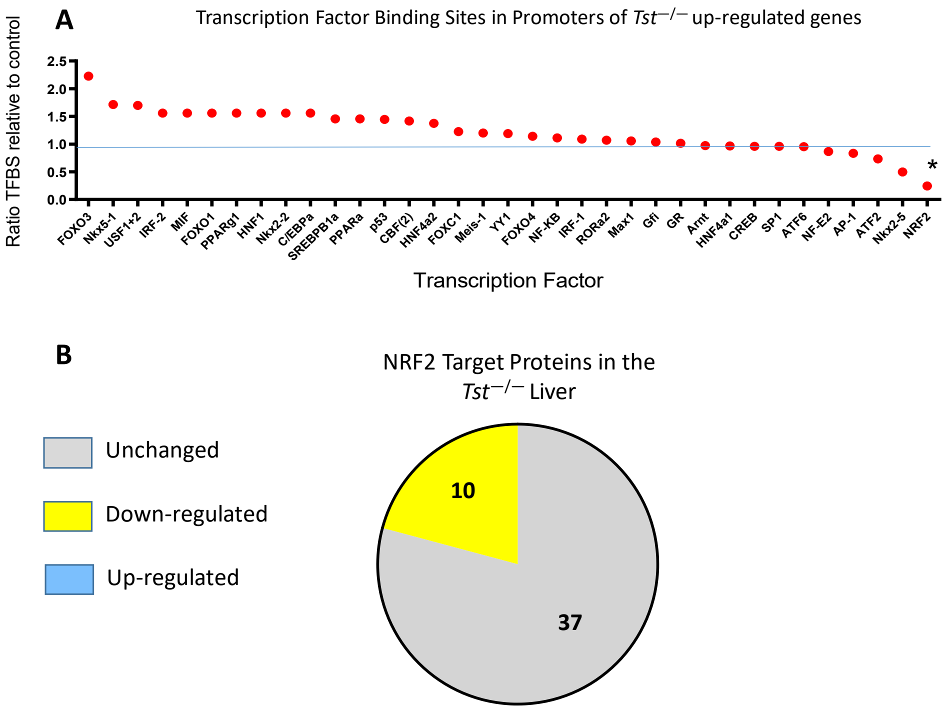


**Figure S6. Hepatic proteins enriched in *Tst* $^{-/-}$ mice show under-representation of NRF2 promoter binding sites.** (A) abundance in *Tst*$^{-/-}$ liver compared to a control set of proteins that are unchanged between 6J and *Tst*$^{-/-}$. The proportion of genes containing a promoter binding site from proteins increased in *Tst*$^{-/-}$ was divided by the proportion of genes containing a binding site from a control set of genes.(B) Pie charts representing the number of NRF2-target proteins whose abundance is increased (blue), decreased (yellow) or unchanged (grey) in the Tst$^{-/-}$ liver. Significance of transcription factor enrichment analysis was calculated using a Fishers Exact test. * $P < 0.05$. Significance for NRF2 target abundance was performed with the Freeman-Halton Fishers Exact Test.

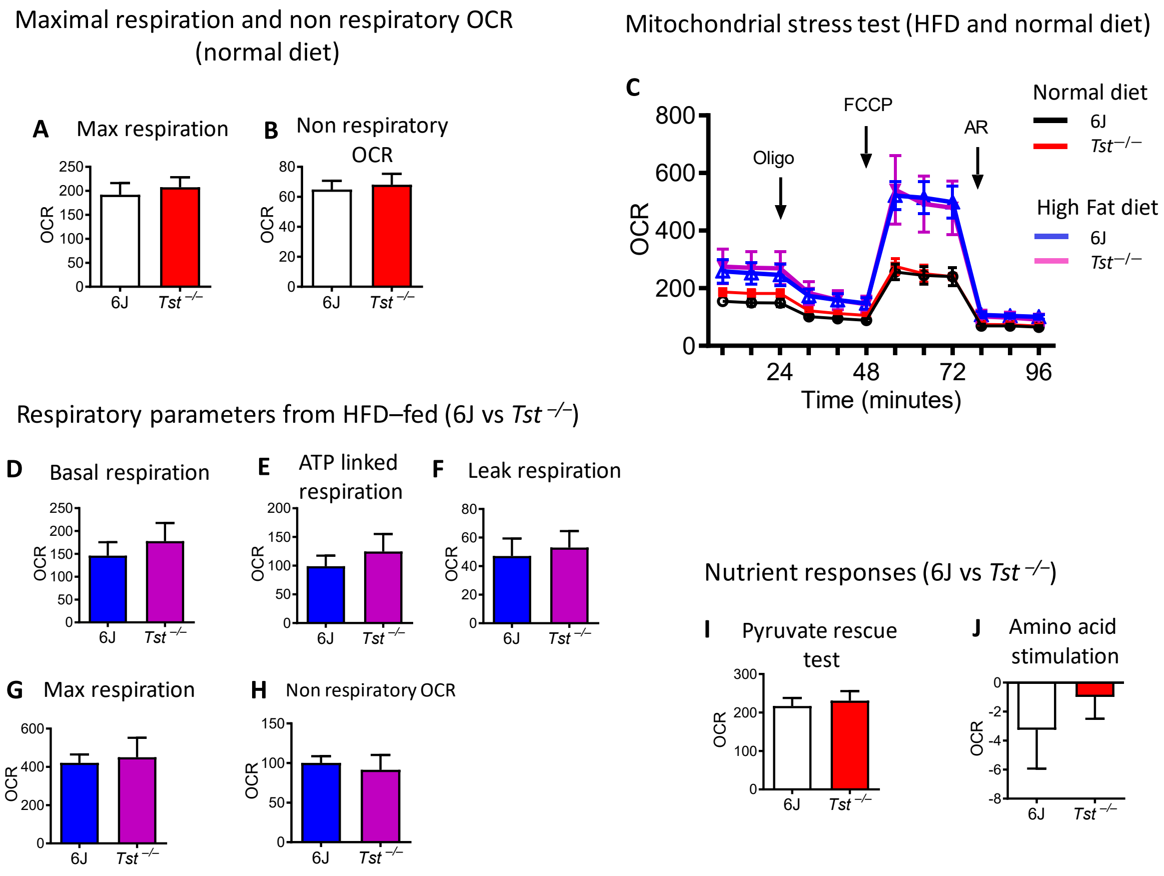


**Figure S7. Hepatocyte respiration after high–fat feeding or after amino acid or pyruvate challenge is comparable between C57Bl/6J and *Tst*$^{-/-}$ mice *in vitro*.** (**A**) Maximal respiratory OCR elicited by uncoupling with FCCP, by hepatocytes from normal diet-fed C57Bl/6J (n = 6) or *Tst*$^{-/-}$ (n = 6) mice, calculated from Figure 3B. (**B**) Non-respiratory OCR remaining following the inhibition of respiration with antimycin and rotenone, by hepatocytes from normal diet-fed C57Bl/6J (n = 6) or *Tst*$^{-/-}$ (n = 6) mice, calculated from Figure 3B. (**C**) Seahorse trace representing the mean oxygen consumption rate (OCR), normalised to protein, by hepatocytes from normal diet-fed (n = 6/genotype), and high fat diet-fed (n = 4/genotype) C57Bl/6J and *Tst*$^{-/-}$mice during a mitochondrial stress test. (**D**)Basal respiratory OCR linked to ATP production (antimycin/rotenone sensitive) by hepatocytes from high fat diet-fed C57Bl/6J (n = 4) or *Tst*$^{-/-}$ (n = 4) mice, calculated from Figure S4C. (**E**) Respiratory OCR linked to ATP production (oligomycin sensitive) by hepatocytes from high fat diet-fed C57Bl/6J (n = 4) or *Tst*$^{-/-}$ (n = 4) mice, calculated from Figure S4C. (**F**) Respiratory OCR relating to proton leak (oligomycin insensitive) by hepatocytes from high fat diet-fed C57Bl/6J (n = 4) or *Tst*$^{-/-}$ (n = 4) mice, calculated from Figure S4C. (**G**) Maximal respiratory OCR elicited by uncoupling with FCCP, by hepatocytes from high fat diet-fed C57Bl/6J (n = 4) or *Tst*$^{-/-}$ (n = 4) mice, calculated from Figure S4C. (**H**) Non-respiratory OCR remaining following the inhibition of respiration with antimycin and rotenone, by hepatocytes high fat diet-fed C57Bl/6J (n = 4) or *Tst*$^{-/-}$ (n = 4) mice, calculated from Figure S4C. (**I**) Stimulation of maximal uncoupled respiration following addition of pyruvate (2mM), from normal diet-fed C57Bl/6J (n = 4) or *Tst*$^{-/-}$ (n = 4) mice. (**J**) Stimulation of maximal uncoupled respiration following addition of aspartate (1mM) and glutamax (1mM), by hepatocytes from normal diet-fed C57Bl/6J (n = 1) or *Tst*$^{-/-}$ (n = 1) mice. Data are represented as mean ±SEM. Significance was calculated using an unpaired two tailed, student's t-test. * $P < 0.05$.

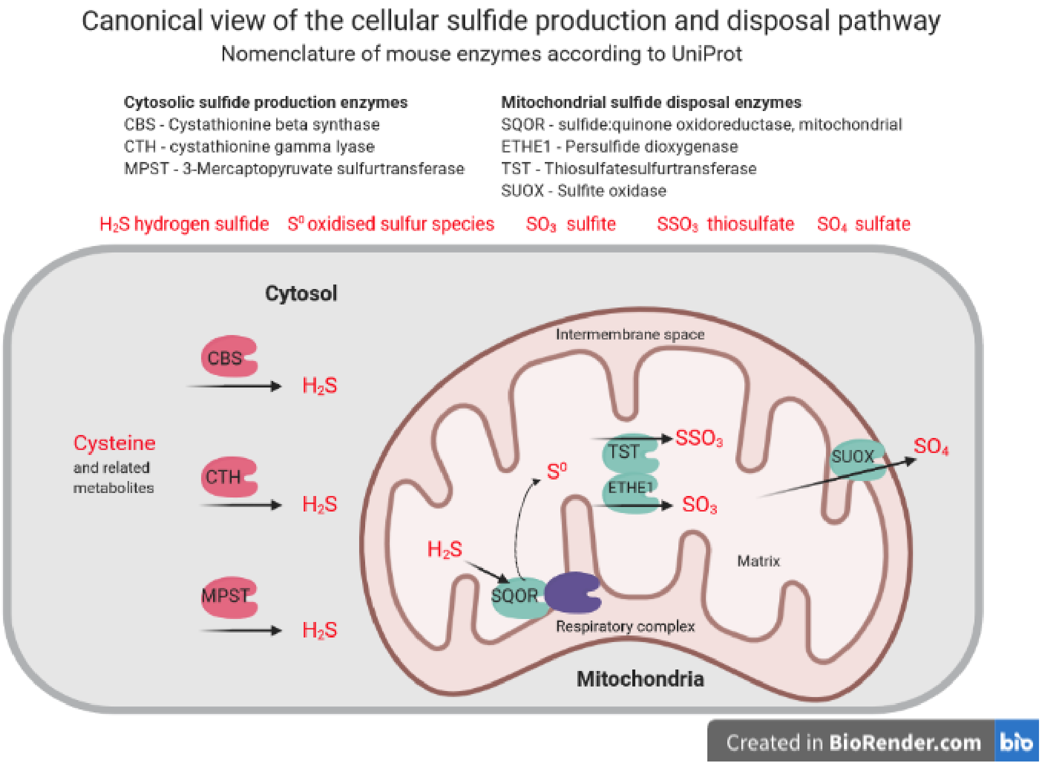


**Figure S8. Canonical view of the cellular sulfide production and disposal pathways.** Intracellular production of $H_2S$ (sulfide) occurs in most cell types (including hepatocytes) by the actions of the cysteine metabolising enzymes in cytosol (CTH, CBS and MPST). The mammalian 'sulfide oxidation pathway', is present in many cell types (including hepatocytes) within mitochondria (SQOR, ETHE1, TST and SUOX). The seven enzymes shown are widely accepted as major contributors to intracellular sulfide metabolism. For simplicity, the diagram does not include sulfide production which can occur within the mitochondria, or disposal pathways in cytosol. The identity of oxidised sulfur species produced at the SQOR and metabolism of other intermediates, including glutathione and glutathione persulfide, remain disputed. The precise role of TST and other enzymes shown here remains under investigation. Nonetheless, the importance of TST 'in vivo' for regulating sulfide metabolism is strongly supported by the sulfide and thiosulfate metabolite measurements we report in the blood of *Tst*$^{-/-}$ mice.

#### Materials and Methods

***Experimental animals.*** All experiments were performed according to guidelines set out by the ethical committees of The University of Edinburgh and Physiogenex S.A.S, Prologue Biotech, Labège, FRANCE. Experiments were carried out within the framework of the Animals (Scientific Procedures) Act (1986) of the United Kingdom Home Office or related laws from the European Union (France). In all studies, animals within genotype cohorts were randomly assigned to diet or intervention groups. All animals were maintained in standard housing with 12 hour light and 12 hour dark cycles (7 a.m. to 7 p.m.) and *ad libitum* access to the appropriate diet. For in vivo experiments (pyruvate tolerance test, insulin tolerance test, euglycaemic clamps), operators and animal handlers were blinded to the data, which was generated by a second individual who was blinded to the treatment regimen until the code was broken. All of the studies used male mice housed in cages of 3-6 individual littermates until intervention. The mice for this study originated from C57Bl/6N $Tst^{-/-}$ mice [1] backcrossed onto the C57Bl/6J genetic background for >10 generations. Mice were placed onto high fat diet D12331, (58% calories from fat, Research Diets, New Brunswick, USA) from between 6-8 weeks of age, for 6-7 weeks prior to testing, and compared to mice maintained on standard low fat diets, RM1 or D12383 (low-fat high-cornstarch, Research Diets, New Brunswick, USA).

***Pyruvate tolerance test.*** Blood glucose was measured from 16 hour fasted mice prior to, and following bolus sodium pyruvate administration (i.p. 1.5 mg/g bodyweight). Blood was collected following tail venesection prior to, and 15, 30, 60 and 120 minutes after injection. Glucose was measured from blood using a Glucometer (*OneTouch*, Lifescan, Milpitas, USA or *Accu-Chek*, Performa nano, Roche).

***PEPCK activity assay.*** Activity of phosphoenolpyruvate carboxykinase was measured from cytosol samples obtained from frozen liver. Samples were homogenised in 250 mM sucrose,

5 mM HEPES, pH 7.4. and centrifuged at 4°C, 12,000 rpm (17,390 g) for 15 min. Supernatants were ultracentrifuged at 4° C, 60,000 rpm (289,000 g) for 30 min. Activity of PEPCK from cytosolic fractions was inferred in this assay from NADH extinction, linked to the conversion of phosphoenol pyruvate into oxaloacetate in the presence of carbonate, dGDP and $MnCl_2$, and the subsequent conversion of oxaloacetate into malate by adding malate dehydrogenase. Baseline measurements at 340 nM (NADH) were taken for 20 min before adding phosphoenol pyruvate, and the reaction proper was initiated with dGDP. The reaction was then measured for a further 40 min.

***Hepatocyte preparations.*** Mice were killed by $CO_2$ asphyxiation, followed by a gentle cervical dislocation. The chest cavity was opened, the portal vein was cut and the thoracic vena cava was cannulated via the right atrium. The liver was perfused with (37°C) perfusion media (140 mM NaCl, 2.6 mM KCl, 0.28 mM $Na_2HPO_4$, 5 mM glucose, 10 mM HEPES, 0.5 mM EGTA, pH 7.4), 6 mls/min for 10 min. The liver was then perfused with digestion media (perfusion media, without EGTA, including 5 mM $CaCl_2$, and 100 U/ml collagenase type 1) for 5-7 min. Finally, the liver was perfused with perfusion media for a further 10 min. Cells were extruded from liver into DMEM medium (DMEM, 5.5 mM glucose, 10% FCS, 7 mM glutamine, and penicillin/streptomycin antibiotics), and then passed through a 40 micron filter. Cells were spun twice and washed with medium, at 500 rpm (47 g) for 5 min. Cells were spun through a 50% Percoll pH 8.5-9.5/DMEM solution at 1000 rpm (190 g) for 15 min to remove dead cells and non hepatocytic liver cell types. Hepatocytes collected in the pellet fractions were resuspended in medium and spun twice with washing at 500 rpm (47 g) 5 minutes. Yields and viability were assessed by counting using a haemocytometer, and proportion of trypan blue exclusion respectively. Yields ranged from between $2 \times 10^6 – 1.5 \times 10^7$ viable cells, and viability was above 85%. Unless otherwise stated, hepatocytes were seeded onto collagen coated tissue culture plastic (collagen from rat tails, Sigma), and maintained in DMEM with 5.5 mM glucose, 10% FCS, 7 mM glutamine, and antibiotics).

***$^{13}C$ Pyruvate metabolite tracing.*** After overnight culture on collagen coated 6-well tissue culture plates, hepatocytes were incubated with 1 mM $^{13}C_3$ labelled pyruvate in serum free DMEM for 60 min. Metabolites were extracted by washing individual wells with ice-cold PBS and addition of cold extraction buffer (50% methanol, 30% acetonitrile, 20% water solution at -20°C or lower). Extracts were clarified and stored at -80°C until required. LC-MS was carried out using a 100 mm x 4.6 mm ZIC-pHILIC column (Merck-Millipore) using a Thermo Ultimate 3000 HPLC inline with a Q Exactive mass spectrometer. A 32 min gradient was developed over the column from 10% buffer A (20 mM ammonium carbonate), 90% buffer B (acetonitrile) to 95% buffer A, 5% buffer B. 10 µl of metabolite extract was applied to the column equilibrated in 5% buffer A, 95% buffer B. Q Exactive data were acquired with polarity switching and standard ESI source and spectrometer settings were applied (typical scan range 75-1050). Metabolites were identified based upon m/z values and retention time matching to standards.

***Plasma lipid analysis***. Mice were fasted with free access to water for 4 hours prior to cull by decapitation or pentobarbital euthanasia. Trunk blood (decapitation) was collected directly into Sarstedt Microvette CB 300 K2E EGTA containing plasma sample tubes (Sarstedt, Nümbrecht, Germany). Venous blood from the abdominal vena cava (post euthanisation) was collected into a BD Plastipak 1 ml syringe (BD, Madrid, Spain). This was then transferred to Sarstedt EGTA containing sample tubes for centrifugation. Blood samples obtained by either method were centrifuged at 20$^{o}$C and 5000 rpm (2655 g) for 5 min to obtain plasma samples. Plasma samples were analysed for cholesterol and triglyceride content by as previously described[2]. Briefly, samples were subjected to gel filtration chromatography using an integrated Alliance HPLC separations module (e2695, Waters, Milford, US) to separate lipoproteins based on size. Effluent was immediately and continuously mixed with either triglyceride (Infinity Triglyceride, Thermo Scientific, Loughborough, UK) or cholesterol (Infinity Cholesterol, Thermo Scientific, Loughborough, UK) enzymatic colourmetric detection kits at the correct conditions for reaction (as specified in manufacturer's guidance). The optical density was then recorded using a spectrophotometer at the appropriate wavelength and the signal turned into a continuous trace i.e. a lipid profile. By identification of the lipoprotein peaks (based on their time of emergence from the chromatograph) the concentration for each could be calculated.

***Oil Red-O lipid analysis of liver.*** 5 µm cryostat cut frozen sections of liver were collected onto Superfrost slides (Thermo), and rinsed with 60% isopropanol. Slides were incubated in freshly prepared staining solution (2.1 mg/ml Oil Red O in 40% isopropanol/water) for 10 – 30 min and rinsed with 60% isopropanol. Slides for representative images were counterstained for nuclei in haematoxylin (Harris) for 1 minute. For image analysis, slides were not counterstained. All slides were then rinsed in running tap water for 2 min, before mounting. Sections were captured using an AxioScan Z1 slide scanner at ×20 magnification and analysis of the proportional area of Oil Red O staining (area of stain/unit area of section) was performed using ImageJ software (National Institutes of Health), assessed by a blinded assessor.

***Liver Glycogen measurement.*** Frozen liver samples (between 30-90 mg) were heated to 100°C in an Eppendorf tube with 0.3 mls of 30% KOH for 30 min with vigorous shaking at 10-min intervals. Samples were heated for a further 2-3 min after addition of 0.1 ml 1M $Na_2SO_4$ and 0.8 ml ethanol. Samples were then centrifuged at 4°C at 1011 g for 5 minutes. The supernatant was removed, and the pellet resuspended in distilled $H_2O$ before 0.1 ml 1M

$Na_2SO_4$ and 0.8 ml ethanol were again added, and samples boiled at 100°C for 5 min before centrifugation. This was repeated a final time to wash the sample. The pellet was resuspended in a 10 mg/ml (~1200 U/ml) amyloglucosidase enzyme in 0.3 M sodium acetate (pH 4.8). Samples were then incubated at 50°C for 2 hours. Quantification of samples was then performed using a standard hexokinase based glucose assay (Glucose (HK) Assay Kit, Sigma, GAHK20). The assay was performed following manufacturer's instructions and values calculated by extrapolation from a standard curve after measuring absorbance using a plate spectrophotometer (Molecular Devices OPTImax microplate reader and software, Molecular Devices, Wokingham, UK).

***Western blotting for protein abundance.*** Frozen liver samples (stored -80°C) from mice were homogenized in protein lysis buffer (50 mM Tris, 270 mM sucrose, 50 mM NaF, 1 mM EDTA, 1 mM EGTA, 1% Triton X-100, 10 mM B-glycerophosphate, 5 mM Na Pyrophosphate, 1 mM orthovanadate, 0.1% β-Mercaptoethanol, 1 tablet protease inhibitor cocktail inhibitor, pH 7.4, all Sigma Aldritch). Samples were then centrifuged at 13,200 rpm (18500 g) for 15 min at 4°C and the supernatants aliquoted and stored at -80°C. Protein samples were loaded onto 10% acrylamide/bis-acrylamide gels (30% acrylamide, Sigma Aldritch) and separated by electrophoresis. A coloured molecular weight marker was also run on all gels (Full range rainbow molecular weight markers, GE Healthcare). Gels were transferred overnight using a Bio Rad wet transfer system onto Amersham Hybond – P membranes (GE Healthcare). After transfer, for normalisation of specific targets to total protein, membranes were stained using the REVERT total protein stain (LICOR), according to manufacturers' instruction. Following stain and wash, lanes of each sample were analysed using a LI-COR Odyssey scanner (700nm channel). For blots using a house keeping protein for normalisation, the total protein stain was not performed, and membranes were transferred directly to blocking. All membranes were blocked in Tris buffered saline with 0.01% tween (TBST, containing 5% skimmed milk powder (Marvel skimmed milk powder) for 1 hour and then rinsed in TBST. Blocked membranes were then incubated with the appropriate primary antibody in TBST containing 5% BSA (Sigma Aldritch) overnight at 4°C. Following three 5 min washes with TBST, secondary antibody incubation for all blots was with an appropriate green or red fluorescent antibody, incubated at room temperature for 2 hours in TBST containing 5% BSA. Membranes were washed three times in TBST then scanned using the LI-COR Odyssey scanner. Odyssey software (LI-COR Biosciences) was used to quantify band intensity. For normalization to a house keeping protein, the individual band intensity of B-actin was used for each sample. Primary antibodies used were; TST, Rabbit, GeneTex, GTX114858, MPST, Rabbit, Abcam, Ab224043, GOT1, Rabbit, Abcam, ab170950, GSTT1, Rabbit, Proteintech, 15838-1-AP, MAT1A, Rabbit, Abcam, ab129176, BHMT Rabbit, Proteintech, 15965-1-AP, CSAD, Rabbit, Abcam, ab91016, PPCS Rabbit, Atlas Antibodies, HPA031361. Secondary antibodies used were; IRDye800CW Goat anti-Rabbit, Li-Cor, 926-32211, IRDye 680RD Donkey anti-Mouse, Li-Cor, 926-68072. For normalisation B-Actin, Mouse, Abcam, ab8226 was used for whole tissue, and Cox IV (mitochondrial loading control), Abcam, ab16056 was used for mitochondrial fractions.

***Insulin tolerance test*.** Male C57BL/6J or *Tst*$^{-/-}$ mice were maintained on standard chow (RM1). Mice were fasted for 4 hours prior to injection i.p. of insulin (1 mU/g bodyweight, NovoRapid 100U/ml, Novo Nordisk). Tail venesection blood samples were taken prior to, and 15, 30, 60 and 120 minutes post injection. Blood glucose was measured from samples using a Glucometer (Accu-Check, Performa Nano, Roche). Blood glucose was plotted across time to evaluate net glucose accumulation in blood.

***Euglycemic hyperinsulinemic clamps.*** Male C57BL/6J or *Tst*$^{-/-}$ mice were maintained on standard diet (RM1 (E) 801492, SDS) or high fat diet for 6 weeks (58% fat, D12331, Research Diets). Prior to performing the hyperinsulinemic euglycemic clamp an indwelling catheter was placed into the femoral vein under isoflurane anesthesia, sealed under the back skin, and glued onto the top of the skull. Clamps were performed 5-6 days after recovery from catheterization. Mice were fasted 6 hours prior to a basal blood sample was taken for glucose and insulin. Mice then received a bolus of D-[3-3H] glucose (30 µCi) and perfused with 3H-glucose (30 µCi/kg/min at 2 µl/min) for 210 min (which covers the basal phase and hyperinsulinemic clamp). At steady state (60 min after start of perfusion), 5 µl of blood was collected and glycemia measured from tail tip every 10 min over 30 min for $^{3}$H-radioactivity analysis for determination of whole body glucose turnover glycolysis and glycogen synthesis rate in the basal state. 90 min after start of perfusion, the hyperinsulinemic clamp starts by co-perfusion with insulin 8 mU/kg/min for the clamped phase over 120 min. Blood glucose was assessed every 10 minutes, and glucose infusion adjusted until steady state blood glucose (120 mg/dl +/- 10 mg/dl) was achieved. 5 µl of blood was collected from, tail tip every 10 min for $^{3}$H- radioactivity analysis. At 150 min after the start of perfusion, a bolus of $^{14}$C-2-deoxyglucose (25 µCi) was perfused to evaluate tissue specific uptake. At the end of the perfusion (210 min), blood is collected from the retro-orbital sinus to measure plasma insulin and mice sacrificed by i.v. injection of pentobarbital and cervical dislocation. Tissues (Inguinal WAT, Epididymal WAT, Soleus muscle, Extensor digitorum longus muscle, Vastus lateralis muscle, Tibialis anterior muscle, Heart apex, Liver) were removed by dissection and flash frozen in liquid nitrogen (stored -80°C until measured). Tracers were used to calculate various aspects of glucose metabolism [3,4]. Parameters measured or calculated include body weight, glucose infusion rate, whole body turnover, hepatic glucose production, whole body glycolytic rate, whole body glycogen synthesis rate, and tissue glucose utilization.

***MBB derivatization of whole blood and plasma.*** Whole blood was taken after cull of mice, from trunk (following decapitation), or portal vein (following $CO_2$ euthanasia). EDTA-plasma was obtained from trunk blood following decapitation and collected onto ice. Blood for plasma was centrifuged within 15 min of collection for 5 min at 5000 rpm (2655 g) at 4°C. Blood and plasma samples (15-50 µl) were derivatized with monobromobimane by addition of 200 µL of 80 mM EPPS (4-(2-Hydroxyethyl)-1-piperazine propanesulfonic acid, 8 mM DTPA (diethylenetriaminepentaacetic acid) pH 8.0, 50% acetonitrile, 2.3 mM monobromobimane. Reaction vials were capped tightly and vortexed for 1 minute and incubated protected from light at room temperature for 30 min. 1 mL ethyl acetate was added, the tube capped and vortexed for 1 min and incubated protected from light for 10 min. The reaction vials were centrifuged at 1800 rpm (350 g) for 7 min to separate aqueous and organic layers. The organic layer was collected from each extraction, transferred to a 1.5 mL brown glass vial and the solvent was evaporated completely under a nitrogen stream. Acetonitrile (200 µL) was added to each vial, and the solvent was again evaporated to remove any traces of ethyl acetate. Dried MBB-derivatives were stored at -20°C until analysed.

***Fluorometric quantification of MBB-sulfur species.*** MBB-sulfur species (sulfide, thiosulfate, reduced glutathione, and cysteine) in samples was quantified by HPLC separation and detection with a fluorescence detector. The dried MBB derivatives were re-suspended in 50 µL of Buffer A (10 mM tetrabutylammonium phosphate aqueous, 10% methanol, 45 mM acetic acid adjusted to pH 3.4). The entire sample was transferred to an HPLC autosampler vial with a 200 µL glass sample insert, and the vial was closed with a penetrable cap. 20 µL of the sample was injected onto a C8 reverse-phase column (LiChrospher 60 RP-select B 5 µm

4.0 × 125 mm LiChroCART 125-4, Merck KGaA) and a guard column (LiChroCART 10-2, Superspher 60 RP-select B cartridge) on an Ultimate 3000 UHPLC+ focused system (Thermo Scientific). MBB derivatives were eluted with a linear gradient from 10% buffer B (10 mM tetrabutylammonium phosphate in methanol, 10% water, 45 mM acetic acid) to 100% buffer B over 30 min. The eluent was analysed by fluorescence (λex = 380 nm, λem = 480 nm).

***Sulfur metabolite analysis from liver.*** Livers from mice were removed promptly following decapitation (within 2 min), and frozen on powdered dry ice. Frozen tissue was pulverized and derivatized with either 2,4-dinitrofluorobenzene for detecting GSH or monobromobimane for detecting sulfide and thiosulfate as described previously [5–7].

***P3 fluorescence detection of sulfide in hepatocytes.*** Hepatocytes were seeded in glass bottomed, collagen coated wells (0.75 $cm^2$, 12,500 hepatocytes per well) and cultured in DMEM with 5.5 mM glucose, 10% FCS, 4 mM glutamax or 7 mM glutamine, and antibiotics overnight. P3 $H_2S$ reactive probe[8] was added to wells at 10 µM in serum free DMEM for 30 min, prior to gentle washing with Krebs phosphate buffered saline (pH 7.4). Plates were measured using the TECAN fluorescence plate reader, following excitation at 375 nm and detection at 510 nm. No-cell control wells were used for subtracting from the cell containing values. Corrected fluorescence emission data was normalised to protein as estimated by sulforhodamine dye. Briefly, after the run cells were fixed with 10% trichloroacetic acid overnight at 4°C. Cells were washed 9 times with tap water, and air dried. Cells were incubated with 200 µl of 0.4% Sulforhodamine dye/1% acetic acid for 1 hour at room temperature. Stain was removed, and washed 4 times with 1% acetic acid, and then air dried. Stain was dissolved in 200 µl of 10 mM Tris pH 10.5 for 30 min, and 100 µl was measured by colorimetric absorbance spectroscopy at 540 nm. After subtracting a baseline absorbance from blank controls, the absorbance was used to normalise the fluorescence data from each well.

***Quantification of hydrogen sulfide levels using MitoA in vivo exomarker.*** MitoA and MitoN were quantified in mouse blood using LC-MS/MS. Mice received a tail vein IV injection of 50 nM MitoA in 0.9% saline (100 µL). MitoA was given 1.5 hr to distribute into mitochondria. Mice were culled by decapitation 90 minutes after administration. Liver was excised and flash frozen in liquid nitrogen. MitoA and MitoN were extracted from tissue by homogenization of liver (50 mg) enriched with 5 pg d15-MitoN (95% ACN, 210 µL) which was used as an internal standard (IS). Homogenates were centrifuged (16,000 g, 10 min, RT) and the supernatant was transferred to a clean tube and stored on ice. The pellet was re-extracted (95% CAN, 210 µL), spun down again (16,000 g, 10 min, rT) and the supernatants were combined and incubated at 4$^{o}$C for 30 mins. Calibration standards comprise MitoA and MitoN standards ranging from 0.01 to 10 pg in 500 µL 95% ACN. 500 µL of the supernatants and calibration standards were loaded onto an ISOLUTE PLD+ protein and phospholipid removal plate (Biotage, Sweden). Samples and standards were pulled through the plate under vacuum into a 2 mL deep-well 96-well plate. Wells were dried completely at 40$^{o}$C under $N_2$ and resuspended in 100 uL 20% ACN, 0.1% FA. The plate was shaken at (250 rpm, 20 min) to ensure reconstitution. Liquid chromatography-Mass Spectrometry was performed on an I-class Acquity LC system-Xevo TQS triple quadrupole mass spectrometer (Waters, Warrington, UK). Samples were kept at 10$^{o}$C and injected onto an Acquity UPLC BEH C18 column fitted with a 0.2 µm filter (1 x 50 mm, 1.7 µm, Waters). Chromatographic separation of MitoA and MitoN was achieved using mobile phase A composition: water:ACN, (95:5, 0.1% FA), mobile phase B: ACN:water (90:10, 0.1% FA). LC mobile phases were infused at 200 µL/min using the gradient: 0– 0.3 min, 5% B; 0.3–3 min, 5–100% B; 3– 4 min, 100% B, 4.0– 4.10, 100–5% B; 4.10– 4.60 min, 5% B. MS/MS analysis was performed under positive ion mode (Source spray voltage, 3.2 kV; cone voltage, 125 V; ion source temperature, 100 °C). Curtain and collision gas were nitrogen and argon, respectively. Analytes were detected by multiple reaction monitoring (MRM). MitoA undergoes neutral loss of N2 to a nitrene (precursor ion). For quantification the following transitions were used: MitoA, *m/z* 437.2 → 183.1; MitoN, *m/z* 439.2 → 120.0; d15-MitoN, 454.2 → 177.1 m/z. MassLynx 4.1 software was used to integrate the peak area of the analytes MitoA, MitoN and the d15-MitoN internal standard. Response was calculated by normalizing sample peak areas to the IS peak area. By comparison of sample responses to calibration standard responses the mass of each analyte in the tissue sample was calculated. The mass of analyte was normalised to the mass of tissue homogenizer and MitoN/MitoA ratio was calculated.

***Preparation of hepatic mitochondria.*** Fresh liver was taken from mice, and homogenised in 250 mM sucrose, 10 mM HEPES, 1 mM EGTA. 0.5% fatty acid free bovine serum albumin (BSA) pH 7.4 at 4°C, with seven passes of a loose glass Dounce homogeniser (Type A). Homogenates were centrifuged in glass tubes at 2900 rpm (1000 g) for 10 min in a pre-chilled 4°C Beckman centrifuge (JA-20 Fixed angle rotor). The supernatant was then centrifuged in glass tubes at 8500 rpm (8700 g) for 10 min at 4°C. The supernatant was aspirated and any visible lipid was carefully removed from the sides of the tubes. The pellet was washed with 5 ml of mIR-05 buffer (0.5 mM EGTA, 3 mM $MgCl_2$, 20 mM taurine, 10 mM $KH_2PO_4$, 20 mM HEPES, 110 mM sucrose, 1 mg/ml fatty acid free BSA, pH 7.2), and centrifuged at 8500 rpm (8700 g) for 10 min at 4°C. After aspiration and removal of visible lipid, the pellet was suspended in 1ml of mIR-05 buffer and kept on ice until used. All measurements were taken within two hours of preparation. Protein concentration was determined using the DC-Protein Assay (BioRad) as per manufacturers instruction.

***Amperometric analysis of sulfide disposal.*** Hepatocytes were prepared as described, and kept at room temperature in DMEM with 5.5 mM glucose, 10% FCS, 4 mM glutamax or 7 mM glutamine, and antibiotics at a concentration of 4 x $10^6$ per ml. Mitochondria were prepared as described, and maintained on ice in mIR-05 buffer until use. All samples were analysed within 4 hours of preparation. Sulfide ($H_2S_{(g)}$) was measured (with and without samples) in a 2ml volume plastic chamber, to which an amperometric sensor was inserted, sealed with a rubber O-ring. Voltage measurements from the sensor (linear relationship to $H_2S_{(g)}$ concentration) were recorded using a TBR4100 Gas radical analyser (World Precision Instruments). A gas permeable membrane covered the sensor, and the outer glass sensor compartment was filled with $H_2S$ detection fluid (World Precision Instruments). All measurements of $H_2S_{(g)}$ from standards and samples were recorded as voltage by the amperometric sensor at ambient temperature. Mitochondrial measurements (and standards) were taken in serum free mIR-05 buffer. Hepatocyte measurements (and standards) were taken in serum-free, bicarbonate-free DMEM, buffered with 25 mM HEPES (pH 7.4), with 5 mM glucose, 2 mM glutamax and 2 mM pyruvate. Sulfide was added to buffer in the form of $Na_2S$, predicted to equilibriate according to its Pka at this pH to about 1/3 of sulfide as $H_2S_{(g)}$ 2/3 as $HS^-$. The probes selectivity to $H_2S_{(g)}$ (vs $HS^-$) was confirmed with standards by demonstrating predicted signal amplification to a maximum following acidification of media to pH < 5 (approx. 100% $H_2S_{(g)}$/0% $HS^-$), and signal compression to a minimum following alkalinisation of standard to pH > 10 (Approx 0% $H_2S_{(g)}$/100% $HS^-/S^{2-}$). A final re-acidification recovered the signal to near maximal levels. Standard curves for calculating experimental measurements were prepared using freshly made $Na_2S$ solutions ranging from 0.25 – 20 µM (corresponding to approximately 170 nm – 6.7 µM $H_2S_{(g)}$). $H_2S_{(g)}$ disposal was measured by recording the extrapolated $H_2S_{(g)}$ concentration after 10 min incubation with samples. A baseline without sample was taken for 5 min, and then after sample addition (400,000 hepatocytes, or 1.6 – 2.0 mg of mitochondrial prep), another 5 min baseline with sample was taken. In all experiments, no detectable increase in signal (limit of detection 0.25 µM $Na_2S$) was observed during incubation of hepatocyte or mitochondrial samples from either genotype. Following addition of 10 µM $Na_2S$ the (voltage) signal was recorded over a period of 10 min. Disposal rates were calculated after subtraction of a baseline disposal rate in media alone, over a 10 min period, performed each day of experimentation. Sample disposal rates were in the range of 5-20 higher than baseline disposal rate confirming good signal to noise. To determine the rate of disposal that is dependent upon respiration, a fresh aliquot of the same sample was prepared as before, but 5 min after addition of sample to chamber, Antimycin A (2.5 µM, dose titrated) was added. After a further 5 minutes, 10 µM $Na_2S$ was added and a disposal rate (after subtraction to sample free baseline rate) was again calculated. The respiratory (Antimycin sensitive/complex III dependent) sulfide disposal rate of samples was calculated as the difference between the nieve sample rate and the Antimycin inhibited rate. After each measurement, the sample was removed, and centrifuged to collect cells or mitochondria for a final protein assessment (DC-Protein Assay, Bio-Rad) for the purposes of normalisation.

***Mitochondrial ROS (MitoSOX) measurement in $H_2O_2$ treated hepatocytes.*** Hepatocytes were seeded overnight onto 96-well collagen coated plates. Cells were exposed to a range of concentrations of $H_2O_2$ (0.125 - 8 µM) for 2 hours. Following 3 washes with PBS, cells were incubated with MitoSOX Red mitochondrial superoxide indicator (Thermo Fisher) for 10 mins prior to three further washes. Measurement of fluorescence was carried out in a fluorescence detector plate reader (TECAN), using 510 nm for excitation and 580 nm for emission detection. Data from each well was normalised to sulfurhodamine dye protein stain.

***Persulfidation Mass Spec and GO term analysis.*** Livers from mice were removed promptly following decapitation (within 2 min), and snap frozen in liquid nitrogen. The persulfide proteome analysis using the BTA method was conducted as described previously[9]. Briefly, 100-150 mg of frozen liver tissue was pulverized and lysed on ice in RIPA buffer (100 mM Tris, pH 7.5, 150 mM NaCl, 2mM EDTA, 1% Triton X-100, 25 mM deoxycholic acid, 2 tablets/ 100 ml of cOmplete™, Mini, EDTA-free Protease Inhibitor Cocktail (Roche)). The lysates were centrifuged at 14,000 g for 10 min at 4°C and protein concentrations were determined using the Bradford reagent (BioRad). Supernatant containing 6 mg of protein was incubated with 100 µM NEM-biotin (Pierce) for 60 min at room temperature after which the proteins were precipitated with cold acetone (1:4 v/v) for 1 h at -20°C, followed by a centrifugation at 14,000 g for 10 min at 4°C. The precipitated protein was re-suspended in a denaturing buffer containing 7 M urea, 1% SDS, 150 mM NaCl, 100 mM Tris, pH 7.5. Then, the samples were diluted 10-fold with trypsin reaction buffer (1 mM $CaCl_2$, 100 mM Tris pH 7.5) and incubated overnight with sequencing grade modified trypsin (1:50 trypsin:protein) (Promega) at 30 °C. The digestion products were mixed with streptavidin-agarose beads (ThermoScientific) and incubated at 4° overnight, followed by ten washes with the wash buffer (0.1 % SDS, 100 mM Tris, pH 7.5, 600 mM NaCl, 1 mM EDTA, 1% Triton X-100). The streptavidin-agarose bound peptides were incubated with elution buffer (100 mM Tris, pH 7.5, 150 mM NaCl, 1 mM EDTA, 30 mM DTT) for 1 hr at room temperature. Persulfidated peptides were eluted by centrifugation and derivatized with 40 mM iodoacetamide for 2 hrs at room temperature in the dark. The samples were then passed through a desalting column (Pierce). LC-MS/MS analysis was carried out using an LTQ-Orbitrap Elite mass spectrometer (Thermo-Fisher) coupled to an Ultimate 3000 high-performance liquid chromatography system. The alkylated peptides were loaded onto a 75 µm desalting column, C18 reverse phase resin (Dionex), and eluted onto a Dionex 15 cm x 75 µm id Acclaim Pepmap C18, 2 µm, 100 Å reverse-phase chromatography column using a gradient of 2–80% buffer B (5% water, 95% acetonitrile, 0.1% formic acid) in buffer A (0.1% formic acid). The peptides were eluted onto the mass spectrometer at a flow rate of 300 nl/min and the spray voltage was set to 1.9 kV.

***GO enrichment analysis.*** In order to identify differentially persulfidated proteins between the C57Bl/6J and *Tst*$^{-/-}$ samples, we compared the abundances of persulfidated fragments in appropriately treated mass spectrometry data sets to the estimated overall abundance of the corresponding parent proteins in standard label-free quantitation experiments. For each observed persulfidated fragment in each experimental replicate, we calculated the persulfidation rate as the $\log_2$ ratio of the count of that persulfidated fragment to the median count of that fragment across all experimental replicates. The observed counts for the *Tst*$^{-/-}$ replicates were additionally scaled (prior to log transformation) by the ratio of abundances of the parent protein between the C57Bl/6J and *Tst*$^{-/-}$ cells to normalize for differential protein abundance across conditions. For each peptide we then assigned an approximate average $\log_2$ fold change in persulfidation rate between the C57Bl/6J and *Tst*$^{-/-}$ conditions. If a persulfidated peptide was identified in at least two biological replicates of one condition and none in the other, we assigned a $\log_2$ fold change of +/- 5.0 as placeholder values indicating a high confidence change; peptides with only one observation in one condition and none in the other were omitted from our analysis. Having thus obtained estimates for the magnitude of changes in persulfidation rate of each detectable peptide, we then performed gene ontology term enrichment analysis using the estimated $\log_2$ fold changes. We consolidated the peptide-level data to protein-level data by taking the largest magnitude change in persulfidation levels across all peptides from a given protein, and then used the iPAGE program [http://dx.doi.org/10.1016/j.molcel.2009.11.016] to identify GO terms with significant mutual information with the profile of persulfidation rates. Arguments to iPAGE were "—max_p=0.1 –minr=0.3 –ebins=9 –exptype=continuous", indicated that the data were discretized into nine equally populated bins prior to analysis, and that default hypergeometric p-value and information content thresholds were relaxed to maximize sensitivity.

***Focussed analysis of persulfidation in gluconeogenesis proteins.*** The gluconeogenesis pathway was selected for a focussed analysis of the persulfidation rate of all cysteine sites detected in the mass spectrometry data as described above. All peptides from proteins present in the persulfidation data set used for GO enrichment analysis, that are defined by the GO term gluconeogenesis (GO 0006094) were included, these were Pgk1, Gpi1, Fbp1 and Tpi1 (22 peptides). The $\log_2$ rate ratio of persulfidation ($Tst^{-/-}$ /6J) of all of these 22 peptides was compared first to the entire mass spectrometry dataset for $\log_2$ rate ratio of persulfidation (1245 peptides after removal of ambiguous peptides, peptides with a P-diff of 0, and the 22 gluconeogenesis peptides). A Mann-Whitney non parametric T-test was used to detect significance. A second analysis was performed with the gluconeogenesis pathway. For this analysis, all $\log_2$ rate ratio's were given a positive sign to indicate the magnitude of change in the $Tst^{-/-}$ relative to 6J, independent to the direction of change. A Mann-Whitney non parametric T-test was then performed to determine if the magnitude of change in persulfidation in the gluconeogenesis pathway was significantly higher than that of the overall the data set.

***Persulfidation labelling and western blotting from frozen liver.*** 80-120 mg of frozen liver samples were homogenized on ice using a 2 ml glass Dounce homogenizer (Kimble), in 500 µl buffer (7 M urea, 100 mM Tris pH 7.5, 150 mM NaCl, 1 mM EDTA, 1% Triton X-100, 1% deoxycholic acid; supplemented with cOmplete Protease Inhibitor Cocktail (Roche)). Initial disruption of tissue was achieved with three passes, using the loose (Type A) pestle, and after

5 min incubation on ice; complete homogenization was achieved with 9 passes using the tight fit (Type B) pestle. Homogenates were centrifuged at 5000 rpm (2655 g) for 5 min at 4°C. Protein concentrations of supernatants were determined using the *DC* BCA protein assay (Bio-Rad). Protein (6 mg) from each sample, was made up to 1 ml with phosphate buffered saline (pH 8.0). Freshly prepared EZ-link Maleimide PEG Biotin EZ-linker (Thermo Fisher 21902BID), was added to samples to 100 µM, and incubated for 1 hour at room temperature with gentle mixing. Excess maleimide linker was removed from samples by acetone precipitation (3 volumes) at -20°C for 1 h, followed by centrifugation at 12000 rpm (17390 g) for 10 min at 4°C. Protein pellets were washed with ice-cold acetone and then dissolved in 250 µl of 50 mM Tris pH 8.0, 100 mM NaCl, 1 mM EDTA, 1 % SDS. To each sample, 750 µl of RIPA buffer (100 mM Tris pH 7.5, 150 mM NaCl, 1 mM EDTA, 1% Triton X-100, 1% deoxycholic acid) was then added. An aliquot (20 µl) was taken from each sample for estimation of total input protein for normalization (described below). The remainder of the samples were split into duplicates and incubated with gentle mixing, overnight at 4°C with 320 µl of pre-washed streptavidin agarose beads (Thermo Fisher 20347). Beads were washed 10 times with 0.8 ml washing buffer (30 mM Tris pH 7.5, 600 mM NaCl, 1 mM EDTA, 1% Triton X-100, 0.1% SDS), followed by one wash with phosphate buffered saline (pH 7.4). Beads were then centrifuged for 1 min at 1000 rpm (106 g) to dry. Elution of the duplicate samples for western blot analysis was performed by adding 300 µl of elution buffer (30 mM Tris pH 7.5, 150 mM NaCl). For each sample pair, one duplicate was eluted in buffer supplemented with 10 mM DTT and the other without DTT. The beads were incubated with the elution buffer for 1 hour at RT; and centrifuged for 1 min at 1000 rpm (106 g) to collect the eluate. Each eluted sample was concentrated to 20 ul using an Ultra-0.5 Centrifugal Filter Device, 10 K cut-off (Amicon), as per the manufacturer's instructions. Eluted samples, and input protein samples were loaded in their entirety onto

SDS PAGE gels and transferred overnight at 4°C by Western blotting onto PVDF membrane. Total protein from each lane on the membrane was estimated after staining with REVERT total protein stain (LICOR) according to the manufacturer's instructions. Briefly, following overnight transfer and after rinsing the blot with water, the membranes were incubated with REVERT total protein stain for 5 min, and rinsed twice with wash solution. Blots were then imaged in the 700 nm channel with an Odyssey imaging system (LICOR). Each lane was measured for its total integrated fluorescence intensity to obtain an estimate of the total protein in each lane. Measurements from no-DTT eluted sample lanes were subtracted from DTT eluted sample lanes. Similar fluorescence measurements of input total protein lanes were used to normalise the eluted sample measurements, and this was used as a measure of relative protein-persulfidation rate.

***Mass spec analysis of liver protein.*** *Sample preparation;* $Tst^{-/-}$ and wild type (C57Bl/6J) mouse strains were fed either high-fat (58% fat) or normal (low fat) diet. Livers from mice were removed following decapitation, and snap frozen in liquid nitrogen. Four biological replicates from the 4 conditions were used to isolated proteins and performed protein quantitation using iTRAQ 8plex. Liver tissue was homogenized using 1 ml of 8 M urea with HEPES buffer pH 8.0. The protein concentration was determined using the Bio-Rad RC DC protein assay kit (Bio-Rad, Hercules, CA, USA). One hundred micro grams of protein from each of the samples were reduced with THP (Tris(hydroxypropyl)phosphine), alkylated with MMTS (methyl methanethiosulfonate) in 500 mM triethylammonium bicarbonate (TEAB, pH 8.5), trypsin digested and subsequently label with iTRAQ 8plex accordingly to the manufacturer's instructions. *Electrostatic Repulsion-Hydrophilic Interaction Chromatography (ERLIC) Peptide fractionation;* Peptide fractionation was performed using a pH gradient. Labeled peptides were dissolved in 100 µL of buffer A (100 mM formic acid, 25% acetonitrile, pH 3.0), followed by fractionation in a 2.6 × 200 mm, 5 μm, 200 Å PolySulfoethyl A column (Poly LC Inc., Columbia, MD), using an Ultimate 3000 UHPLC+ focused (Thermo-Fisher Scientific) system, operating at a flow rate of 0.2 ml/min. Twenty minutes of isocratic buffer A were followed by a linear gradient from 0% to 100% buffer B (100 mM ammonium formate, 25% acetonitrile, pH 6.0) over 20 min and then a final linear gradient from 0% to 100% buffer C (600 mM ammonium acetate, 25% acetonitrile, pH 6.0) over 10 min. A total of 22 fractions (1-min intervals) were collected. All fractions were lyophilized and stored at -20 °C. *Nanoflow Liquid Chromatography Tandem Mass Spectrometry;* NanoLC MS/MS analysis was performed using an on-line system consisting of a nano-pump UltiMate™ 3000 UHPLC binary HPLC system (Dionex, ThermoFisher) coupled with Q-Exactive mass spectrometer (ThermoFisher, San Jose, CA. iTRAQ-labeled peptides were resuspended in 2% ACN, 0.1% formic acid (20 µL) and 6 µL injected into a pre-column 300 µm×5 mm (Acclaim PepMap, 5 µm particle size). After loading, peptides were eluted to a capillary column 75 µm×50 cm (Acclaim Pepmap, 3 µm particle size). Peptides were eluted into the MS, at a flow rate of 300 nL/min, using a 90 min gradient from 0% to 35% mobile phase B. Mobile phase A was 2.5% acetonitrile with 0.1% formic acid in $H_2O$ and mobile phase B was 90% acetonitrile with 0.025% trifluoroacetic acid and 0.1% formic acid. The mass spectrometer was operated in data-dependent mode, with a single MS scan in the orbitrap (400-2000 m/z at 70 000 resolution at 200 m/z in profile mode); automatic gain control (AGC) was set to accumulate $4 \times 10^5$ ions, with a maximum injection time of 50 ms. MS/MS scans were performed in the orbitrap at 17 500 resolution. Ions selected for MS/MS scan were fragmented using higher energy collision dissociation (HCD) at normalized collision energy of 38% with an isolation window of 0.7 m/z. MS2 spectra were acquired with a fixed first m/z of 100. The intensity threshold for fragmentation was set to 50 000 and included charge states 2+ to 7+. A dynamic exclusion of 60 s was applied with a mass tolerance of 10 ppm. *Data Analysis;* Raw files were converted to MGF files and searched against the mouse UniProt database (81033 sequences, released on March 2014) using MASCOT Version 2.4 (Matrix Science Ltd, UK). Search parameters were peptide mass tolerance of 10 ppm, and MS/MS tolerance of 0.05 amu allowing 2 missed cleavage. iTRAQ8plex (N-term) and iTRAQ8plex (K) were set as fixed modification, and acetyl (Protein N-term), Methylthio (C) and Oxidation (M) were allowed as variable modification. Peptide assignments with ion score cut-off of 20 and a significance threshold of $\rho < 0.05$ were exported to Excel for further analysis.

***GO and KEGG enrichment analysis of proteome data.*** The data generated from the initial mass spectrometric analysis of iTRAQ labelled peptides from the 16 liver samples was analysed by FIOS. A total of 16 samples were QC analysed using the arrayQualityMetrics Bioconductor package to identify sub-standard and/or outlier samples. No samples were identified as outliers. All samples passed the manual and automated quality control based on three metrics (MAplot, Boxplot and Heatmap). The exploratory analysis using PCA showed that the samples clustered perfectly into four groups based on the factor Group (representing four genotype-diet combinations). The first PC captures the main source of variation in the dataset and is showing a separation of the samples based on diet, where high-fat diet and control diet samples separate. The second PC shows a separation between genotypes (Tst KO and WT). The hierarchical clustering and PCA plot both show a clear separation based on the iTRAQ labels. This is expected as the iTRAQ labels are confounded with the Groups. While the observed separation of the samples into groups is most likely due to the underlying biological differences, any technical variations (potentially introduced during the wet lab processing) could be masked. The log2 ratio data were subsequently normalised within arrays using loess, followed by normalisation between samples using the Gquantile method. A total of 4 single and/or multi-factor comparisons, using statistical approaches, were performed. The contrast

"Tst KO vs WT mice (High-fat diet)" was analysed at a cut-off (unadjusted) p-value < 0.01. Due to the known bias in fold-change magnitudes of the iTRAQ technology, no fold-change cut-off was applied to the significant differentially abundant proteins. With this threshold 551 unique proteins were differentially abundant in at least one of the comparisons. The contrast "High-fat diet vs Control diet (WT)" had the most DAPs (432) while the contrast " $Tst^{-/-}$ vs 6J mice (High-fat diet)" had the least DAPs (83). Noticeably, the TST protein showed the strongest down-regulation for both of the contrasts comparing $Tst^{-/-}$ to 6J mice, consistent with gene deficiency and the fold change compression effect of iTRAQ. The full dataset (4,322 identified proteins) was filtered to remove proteins having less than two detected peptides (on average across all 16 samples); leaving 1,654 proteins for downstream analysis. Exploratory analysis using principal component analysis (PCA) showed that the dataset separated into four distinct groups based on the genotype-diet combinations along the two first principal components (PCs). These 1,654 proteins were used for enrichment analysis for GO terms and KEGG pathways. Individual proteins were considered of interest if they were found significantly different (P < 0.01) between selected pairwise comparisons. The four comparisons were $Tst^{-/-}$ normal diet vs C57Bl/6J normal diet, C57Bl/6J high fat diet vs C57Bl/6J normal diet, $Tst^{-/-}$ high fat diet vs $Tst^{-/-}$ normal diet, and $Tst^{-/-}$ high fat diet vs C57B/6J high fat diet. Normalised mean abundance of proteins was expressed as Log2 fold change ratios for each comparison.

***Transcription factor enrichment analysis.*** 43 up-regulated proteins were selected for analysis of their promoter sequences (selected on basis of P-value < 0.05; adjusted for comparison of diet and genotype). 67 control proteins were selected from the proteome data on the basis of their equivalence of abundance between C57Bl/6J and $Tst^{-/-}$. We used a QIAGEN hosted/SABiosciences mouse database (sabiosciences.com) of promoter located transcription factor binding sites. 34 transcription factors were chosen to analyse, based on either their a-priori prevalence in the promoter of *Tst*$^{-/-}$ up-regulated proteins (present in the promoters of more than 50% of the up-regulated proteins) or on their links to either sulfide or nutrient metabolism. The proportion of genes containing a TFBS was calculated for the up-regulated set (43) and the control set (67). The ratio of up-regulated to control was then calculated. The number of genes with and without the presence of the TFBS were analysed for establishing statistical difference (Up-regulated vs Control), using a Fisher Exact test ($P < 0.05$).

***NRF2 target identification and proteome analysis.*** NRF2 target genes of the mouse liver were compiled from the following reviews [10-13]. 106 genes were identified as target genes (upregulated at mRNA or protein level following NRF2 activation). 47 of these target genes were represented in our liver proteome, and each protein was checked for relative expression between *Tst*$^{-/-}$ and 6J (on normal diet, threshold of $P < 0.01$). 10 of the 47 target genes were lower in abundance in the *Tst*$^{-/-}$ proteome, 37 unchanged, with none upregulated. To analyse whether this was statistically significant, we compared this to the percentage of proteins upregulated, down-regulated or unchanged in the proteome database. 5.86% of proteins were upregulated, 5.62% downregulated, and 88.6% unchanged in the full database (1654 proteins total). Expected (mean) numbers of proteins from a hypothetical set of 47 proteins, predict rounded values of 3 upregulated, 3 downregulated and 42 unchanged. We used these as a reference to the actual data for NRF2 target proteins; 0 upregulated, 10 downregulated and 37 unchanged. A Freeman-Halton Fisher Exact test was used for analysis of significance, and a significant difference between predicted and actual distribution was found ($P_A = 0.039$, $P_B = 0.047$).

***Electron micrograph imaging.*** Liver tissue for transmission electron microscopy was prepared following immersion fixation in 0.1 M PB buffer (pH 7.4, EM-grade) containing 4% paraformaldehyde and 2.5% glutaraldehyde. 1mm tissue blocks were post-fixed in 1% osmium tetroxide in 0.1 M PB for 45 min before dehydration through an ascending series of ethanol solutions and propylene oxide. Tissue blocks were then embedded in Durcupan before ultrathin sections (~60/70 nm) were cut and collected on formvar-coated grids (Agar Scientific, UK), stained with uranyl acetate and lead citrate in an LKB Ultrostainer and then quantitatively assessed in a Philips CM12 transmission electron microscope (TEM).

***Seahorse respiratory analysis.*** Primary hepatocytes (C57Bl/6J and $Tst^{-/-}$ mice) were seeded immediately following purification onto collagen coated V7 Seahorse 24-well cell culture microplates (Agilent Technologies), in 200 µl medium (DMEM, 5.5 mM glucose, 10% FCS, 7 mM glutamine, and penicillin/streptomycin antibiotics), for culture in a 5% $CO_2$ 37°C incubator. Experiments were performed between 22-28 hours following collection from mice. Optimisation experiments determined the optimal seeding density, which was then standardised at 10,000/well. Optimisation for drugs and compounds used in Seahorse experiments were performed separately with hepatocytes for each genotype and dietary regime (normal or 58% high fat). This established the doses of drugs for respiratory manipulation, which were the same for both genotypes and diets; oligomycin (2 µM), FCCP (0.5 µM), and antimycin/rotenone (1 µM/0.2 µM). In all experiments, overnight media from cells was replaced, after two washes (0.75 ml), with 525 µl of run media and incubated for 30 mins at 37°C (without $CO_2$), prior to entry into the Seahorse XFe24 Extracellular Flux Analyser (Agilent). The analyser was operated using Wave software (Agilent), and all oxygen consumption rate (OCR) data was normalised to protein using the Sulforhodamine B stain (described above). Data from each biological replicate was averaged from between 4-10 replicate wells, to produce a single value at each measurement time for each biological replicate. Respiratory parameters were calculated as described below for each biological replicate, and this data was used for statistical analysis of genotype effects.

***Mitochondrial stress test (MST).*** Run media for the MST was Seahorse assay media (Agilent), supplemented with 10 mM glucose, 2 mM sodium pyruvate, pH 7.35 ± 0.5 at 37°C). Most measurements were made using 3 min mixing, 2 min wait, 3 min measure. Measurements following addition of FCCP to hepatocytes from high fat fed mice were measured using 4 min mix, 2 min wait, 2 min measure. Three measurements were taken basally, and three measurements taken after injection of each drug (in sequence; oligomycin for inhibiting ATP-linked respiration, FCCP for eliciting maximal uncoupled respiration, antimycin/rotenone for inhibiting the respiratory electron chain). Respiratory parameters for each biological replicate were calculated from the mean normalised OCR as follows. *Basal respiration* was calculated by subtracting the third OCR measurement following injection of antimycin/rotenone (12$^{th}$ measurement) from the third basal OCR measurement (3$^{rd}$ measurement). *ATP linked respiration* was calculated by subtracting the third OCR measurement following the injection of oligomycin (6$^{th}$ measurement) from the third basal OCR measurement (3$^{rd}$ measurement). *Maximum (uncoupled) respiration* was calculated by subtracting the third OCR measurement after injection of antimycin/rotenone (12$^{th}$ measurement) from the first measurement (peak OCR) following injection of FCCP (7$^{th}$ measurement). *Proton leak respiration* was calculated by subtracting the third OCR measurement after injection of antimycin/rotenone (12$^{th}$ measurement) from the third measurement following the injection of oligomycin (6$^{th}$ measurement). *Non-respiratory OCR* was taken from the third measurement after the addition of antimycin/rotenone (12$^{th}$ measurement).

***Octanoate rescue test.*** To investigate lipid respiratory metabolism, hepatocytes were prepared, seeded and cultured overnight as above. Run media for the Octanoate rescue was Seahorse assay media (Agilent), supplemented with 5 mM glucose, 0.1 mM sodium pyruvate, 1 mM sodium lactate, and 0.5 mM carnitine pH 7.35 ± 0.5 at 37°C). All measurements were made using 3 min mixing, 2 min wait, 3 min measure. After washing cells into run media, and 30 min before entry into the analyser, half of the wells from each genotype were incubated with 8 µM etomoxir (or DMSO vehicle) to block carnitine dependent import of long chain fatty acids into the mitochondria. Three basal measurements were taken prior to injection of oligomycin, two measurements were taken prior to FCCP, two measurements were taken prior to injection of sodium octanoate (250 µM), three measurements taken prior to antimycin/rotenone followed by two final measurements. Standard respiratory parameters were calculated analogous to the above description for the standard mitochondrial stress test, except using the second measurement following injection of drug when only 2 measurements were taken. Dependency upon endogenous fatty acids for supporting uncoupled respiration (*Etomoxir inhibited respiration*) was calculated for each biological replicate using the maximal respiration prior to the addition of octanoate. Maximal respiration was calculated as the 6$^{th}$ measurement (peak FCCP OCR) – 12$^{th}$ measurement (lowest Antimycin/Rotenone OCR). The mean maximal respiration from the etomoxir treated wells was subtracted from the mean maximal respiration of the vehicle treated wells to calculate the Etomoxir inhibited respiration (long chain fatty acid dependency) for that biological replicate. Octanoate stimulation of respiration (*Octanoate stimulated respiration*), was calculated for each vehicle well by subtracting the second OCR measurement after injection of FCCP (7$^{th}$ measurement) from the third measurement after injection of octanoate (10$^{th}$ measurement).

***Real time for mRNA analysis.*** RNA extraction, cDNA synthesis and real-time PCR were performed as described [14,15]. Probes were mouse *Mpst*, Mm00460389_m1, *Tst*, Mm00726109_m1; *Gapdh* (internal control), Mm99999915_g1; and *Tbp* (internal control), Mm0000446973_m1.

***Quantification and Statistical Analysis.*** For analytes, bioenergetics, fluorescent probes, gene expression, and protein levels, generally group sizes of 6 were calculated to allow detection of differences in these variable parameters to a threshold of 15% (there is sufficient power to detect smaller differences in certain parameters with this cohort size) with a power of at least 0.8. In some studies, limitations in animal numbers, or fewer remaining samples from larger group sizes resulting from their use for multiple end-points, precluded the desired minimum of $n$ = 6 per group. Protein or mRNA differences in validation studies with 2 parameters (e.g. diet with line or genotype) were analysed using 2-way ANOVA for line and diet effects followed, where appropriate, by post-hoc Tukey tests or Holm-Sidak multiple comparison tests using Sigmastat version 3.5 (Systat Software) or Prism (Graphpad Software). For simple 2 condition comparisons, *t*-test was used. For simple control versus treated (including different treatments or concentration response curves) data were analysed by 1-way ANOVA. For longitudinal measures (e.g. PTT, ITT, bodyweight gain) repeated measures (RM) ANOVA was used and multiple comparisons determined. For all main *in vivo* studies, a blinding strategy was used where the operator (e.g. for injections of glucose, or administration of drug) was blind to the genotype of the subject during the experiment. Similarly, for analysis of images (e.g. oil-red O staining) the scorer was blind to genotype and the data coded, with the code broken by a second individual. Downstream analysis of e.g. tissue mRNA and protein content was not generally blinded to allow appropriate data arrangement on e.g. representative western blots. For clamp studies, mean ± standard error of mean (sem) will be presented, statistical analysis will use t-test to investigate differences of genotype on each diet (2 independent experiments, normal diet, and high fat diet, are not compared directly to each other).

15. Moreno-Navarrete JM, Petrov P, Serrano M, et al. Decreased RB1 mRNA, protein, and

activity reflect obesity-induced altered adipogenic capacity in human adipose tissue. *Diabetes*. 2013;62(6):1923-1931. doi:10.2337/db12-0977
